# *Acorde*: unraveling functionally-interpretable networks of isoform co-usage from single cell data

**DOI:** 10.1101/2021.05.07.441841

**Authors:** Angeles Arzalluz-Luque, Pedro Salguero, Sonia Tarazona, Ana Conesa

## Abstract

Alternative splicing (AS) is a highly-regulated post-transcriptional mechanism known to modulate isoform expression within genes and contribute to cell-type identity. However, the extent to which alternative isoforms establish co-expression networks that may relevant in cellular function has not been explored yet. Here, we present *acorde*, a pipeline that successfully leverages bulk long reads and single-cell data to confidently detect alternative isoform co-expression relationships. To achieve this, we developed and validated percentile correlations, a novel approach that overcomes data sparsity and yields accurate co-expression estimates from single-cell data. Next, *acorde* uses correlations to cluster co-expressed isoforms into a network, unraveling cell type-specific alternative isoform usage patterns. By selecting same-gene isoforms between these clusters, we subsequently detect and characterize genes with co-differential isoform usage (coDIU) across neural cell types. Finally, we predict functional elements from long read-defined isoforms and provide insight into biological processes, motifs and domains potentially controlled by the coordination of post-transcriptional regulation.

## Introduction

Single-cell RNA-seq (scRNA-seq) has revolutionized transcriptomics analysis, especially as the development of technologies with increasingly higher throughputs has enabled the processing of thousands of single cells simultaneously, boosting the amount of biological diversity that can be captured in a sequencing experiment^1^. The technology has been extensively applied to the discovery of new cell types and the characterization of their transcriptional profiles, resulting in the definition of cell type marker genes^2–7^. Even though scRNA-seq datasets can encode several levels of granularity in the form of cell subtypes, these studies rely on the low number of features required to recapitulate the cell type structure of the data^8^, which situates cell type characterization efforts at the baseline of understanding the intricacy of single-cell biology. scRNA-seq studies have also tackled transcriptional dynamics and how they relate to cell type properties. These include methods for the study of dynamic processes, namely pseudotime^9, 10^ and RNA-velocity^11^ analyses, which have provided insight on cell differentiation and transition mechanisms between cell states^7, 12–16^. Moreover, novel methods have been developed to convey the inference of gene regulatory networks (GRNs) to the single-cell level^17^ in an attempt to combine single-cell information with extant knowledge to infer relationships between genes and transcriptional regulators at a higher resolution.

Single-cell research is nevertheless far from realizing its full potential. On the contrary, the timing is now optimal for the field to undertake the investigation of deeper layers of cellular complexity. In particular, the investigation of Alternative Splicing (AS) and isoform expression dynamics has remained a challenge to the field. Reasons for this include the uncertainty of short read-based isoform quantification, which is exacerbated by the lower number of available reads per transcript in comparison to bulk libraries^18^, and the fact that the most popular scRNA-seq methods are heavily 3’ end biased, which precludes the unambiguous identification of alternative transcript variants^19^. Current methods for the study of isoforms in single cells therefore rely on alternative metrics that either avoid isoform-level expression estimation completely^20, 21^ or exclusively consider individual splicing events^22–26^, generally leaving isoform characterization aside, with some recent exceptions^27^. Meanwhile, long read RNA sequencing (lrRNA-seq) of single cells is beginning to emerge an alternative approach to mitigate this ambiguity, given that it successfully grasps how individual events are combined into alternative isoforms^19^. Long read studies have therefore expanded the field’s notions of cell type-specific splicing from event inclusion towards isoform selection patterns and showed that cell type-specific isoform expression can be detected in both broad types as well as cell subtypes^28–32^. Unfortunately, the sequencing depth constraints intrinsic to long read protocols^19^ have limited the amount of isoform diversity that can be captured by single-cell long read transcriptomics^28–30^, and datasets generally show low levels of redundancy between cells of the same cell type.

Notwithstanding this limited scenario, there are a number of relevant questions regarding the importance of splicing for cell identity and function that can only be resolved by evaluating isoform expression at the single-cell level. In point of fact, splicing differences have been shown to discriminate cell types with an accuracy comparable to that obtained using gene expression^33^, while integrating AS and gene expression changes has led to the discovery of cell subtypes and states that were otherwise not detected^27, 34–36^. Especially relevant among these inquiries is the much-debated issue of whether individual cells express one or several isoforms, that is, whether the isoform diversity observed in bulk studies is recapitulated by each single cell or, alternatively, arises as a result of the combination of multiple cells, each of which uniquely express one of the gene’s isoforms. Ever since the publication of the first scRNA-seq studies, short reads have been used to answer this question, usually via the characterization of splicing event -rather than isoform expression-modalities. Successive studies have provided non-conclusive results, with evidence of bimodal splicing patterns^22, 37, 38^ as well as concerns regarding the relationship between bimodal isoform detection and technical noise^18, 39^, a controversy that suggests that new analytical and computational approaches are needed to understand the isoform landscape of single cells.

Another pending question for the field is whether isoform expression programs involve co-expression relationships between transcript variants from different genes. So far, the application of long read technologies to single-cell data has served to unravel coordinated event choice patterns within isoforms of the same gene^40, 41^, however, cross-gene isoform expression networks have not been investigated. In other words, there have been no studies addressing potential codependency between genes regarding the selection of transcript variants from their isoform repertoire, or the implications of this coordination for cell-state and cell-type properties. This is not only related to the general constraints of single-cell isoform studies, but also to the lack of computational methods and mathematical models to extract this complex signal from the data. In spite of the present research gap, isoform co-expression networks are an anticipated consequence of the regulation of splicing by RNA binding proteins (RBPs) and other splicing factors, and their investigation constitutes an opportunity to gain insight on the functional role of AS. Moreover, given that a multiple cell type and high cell throughput context such as that of scRNA-seq data constitutes a far more suitable data scenario to that of bulk RNA-seq, this is undoubtedly a timely inquiry to make.

In the present study, we hypothesize that isoform expression coordination exists as a result of AS regulation, and that this can be computationally detected in the form of isoform groups showing co-variation across cell types. To demonstrate this hypothesis, we have designed an end-to-end, data-intensive pipeline for the study of isoform networks (Figure 1). First and foremost, we employed a hybrid strategy where bulk long-reads and single-cell Illumina sequencing were integrated to estimate isoform expression at the single cell level. To unlock the limitations of extant correlation metrics in the single-cell context^42^, we developed a novel strategy to obtain noise-robust correlation estimates in scRNA-seq data, and a semi-automated clustering approach to detect modules of co-expressed isoforms across cell types. We additionally re-defined and implemented Differential Isoform Usage (DIU) and co-Differential Isoform Usage (coDIU) analyses in order to leverage the multiple cell types contained in single-cell datasets. Finally, to couple these analyses with a biologically interpretable readout, we incorporated a functional annotation step in which several databases and prediction tools were integrated to add isoform-specific functional information (Figure 1).

**Figure 1:**
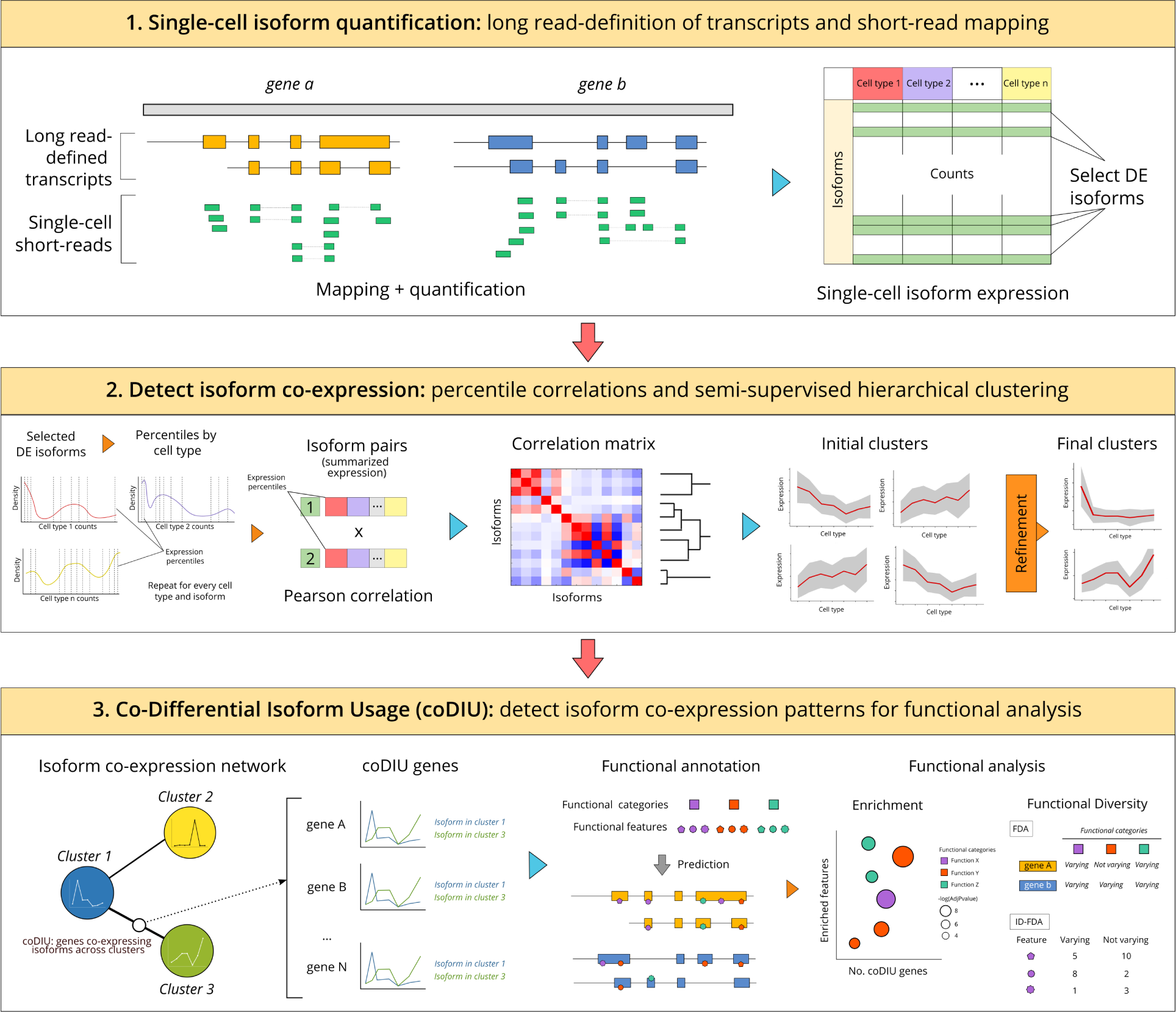
acorde workflow. The *acorde* pipeline includes three main analysis modules. First, long read RNA-Seq da-ta is used to define isoform models and short, single-cell RNA-Seq reads are mapped to the long read-generated transcriptome. Isoform are then tested for multi-group differential expression and those that are significantly DE in at least one of the cell types are selected. Next, percentile correlations are computed to cluster isoforms with similar expression patterns across cell types. Finally, gene pairs are tested for co-differential isoform usage, detecting genes that form co-expression relationships for subsequent functional analysis.

We have hereby applied this pipeline (Figure 1) to the analysis of a publicly available mouse neural dataset, including published scRNA-seq Smart-Seq2 (primary visual cortex, generated by Tasic *et al.*^43^) and bulk ENCODE PacBio long-read data (mouse cortex and hippocampus, generated by Wyman *et al.*^44^). As a result, we successfully detected cell type-specific co-expression of isoforms in a manner that was independent of gene-level expression.

Furthermore, we demonstrated that these isoforms encode shared functional properties, highlighting the role of post-transcriptional processing as an additional regulatory layer that fine-tunes cellular functions and contributes to encode cell type identity. This novel pipeline has been implemented in the R package *acorde* (https://github.com/ConesaLab/acorde).

## Results

### Enabling multi-group differential expression of isoforms in single-cell data

Traditionally, RNA-seq studies use publicly available reference transcriptomes such as RefSeq and ENSEMBL for short read isoform quantification. However, most tissues and cell types will express only a subset of the genes and isoforms contained in the reference, with previous studies showing that isoform detection accuracy increases when adopting tissue-specific isoform sets as a reference for mapping^45^. Long read technologies have the potential to achieve this, while also expanding the reference with novel isoforms that have not yet been annotated^46^. Given the low depth of single-cell long read datasets^19^, we employed a bulk PacBio dataset obtained from ENCODE^44^ to define a mouse, neural-specific transcriptome, that, after extensive curation using the SQANTI3 toolkit (https://github.com/ConesaLab/SQANTI3) contained 36,986 isoforms belonging to 12,692 genes (see Supplementary Note 1).

To quantify the expression of the long read-defined isoforms at the single-cell level, we made use of a publicly available, deeply sequenced, full-length, short-read single-cell RNA-seq dataset by Tasic *et al.*^43^. We retained 1,591 cells after quality control (see Methods). Using the labels from the original characterization of the dataset, we assigned cells to 7 broad cell types, 5 glial (microglia, endothelial cells, oligodendrocytes, oligodendrocyte precursor cells (OPCs) and astrocytes) and 2 neural (GABA-ergic and glutamatergic neurons), each of which can be divided into several, distinct subtypes (Supplementary Figure 1A). To target potential differential isoform selection between cell types, we selected genes that retained more than one isoform after quality filtering, ultimately keeping 13,832 isoforms from 4,591 genes.

Of note, we observed a drastic cell number imbalance between neural (∼720 cells/cell type) and glial cell types (∼30 cells/cell type) in the Tasic dataset, which resulted in the underestimation of transcriptional differences between non-neural types (Supplementary Figure 1B). In order to facilitate downstream analysis, we used a downsampling approach (Methods) to balance the cell type abundances in the dataset while effectively preserving the data structure (Supplementary Figure 1C). Next, to select isoforms with robust co-variation and non-constitutive expression, we applied a multi-group strategy to detect isoforms showing Differential Expression (DE) in at least one cell type, combining the ZinBWaVE zero-expression weighting strategy^47^ with bulk-designed DE methods DESeq2^48^ and edgeR^49^ (see Methods). Upon testing, we detected 5,711 DE isoforms using edgeR (FDR<0.05) and 7,714 using DESeq2 (FDR<0.05). To maximize sensitivity and ensure a broad iso-transcriptome analysis, we considered an isoform to be DE if it was detected by at least one of these methods, resulting in 10,100 isoforms from 4,305 genes (Supplementary Figure 2). Since two or more isoforms with differential cell type expression are required to form co-splicing relationships (see Methods), we finally retained 8,967 isoforms from 3,172 genes with multiple DE isoforms for downstream analysis.

To ensure that downsampling had no effect on DE results, a validation experiment for multi-group DE analysis was conducted, in which we performed 50 runs of random neural cell sampling (see Methods). Although DESeq2 proved to be slightly more robust across independent runs than edgeR, Jaccard Index values indicated that the majority of isoforms were consistently detected as DE (DESeq2: mean no. of DE isoforms = 6,908±101, mean Jaccard Index = 0.84±0.02; edgeR: mean no. of DE isoforms = 6,016±410, mean Jaccard Index = 0.74±0.02). In addition, this validation run showed that considering the union of edgeR and DESeq2 results contributed to improve robustness (mean no. of DE isoforms in union between methods = 9,399±248, Jaccard Index = 0.82±0.01).

### Detecting isoforms showing cell type-level co-expression

Co-expression signals in single-cell data are weak and often result in poor performance of traditional correlation metrics and network inference methods^42, 50^. Although data transformation approaches^51^ and alternative metrics^42^ have been proposed, these are more complex to apply and considerably less interpretable, respectively. Furthermore, most of these studies have only investigated gene level co-expression^17^, often ignoring the AS regulatory landscape. To address these limitations, we implemented a *percentile correlation* strategy: a simple, scalable approach to overcome single-cell noise in isoform co-expression studies (Figure 2A).

**Figure 2:**
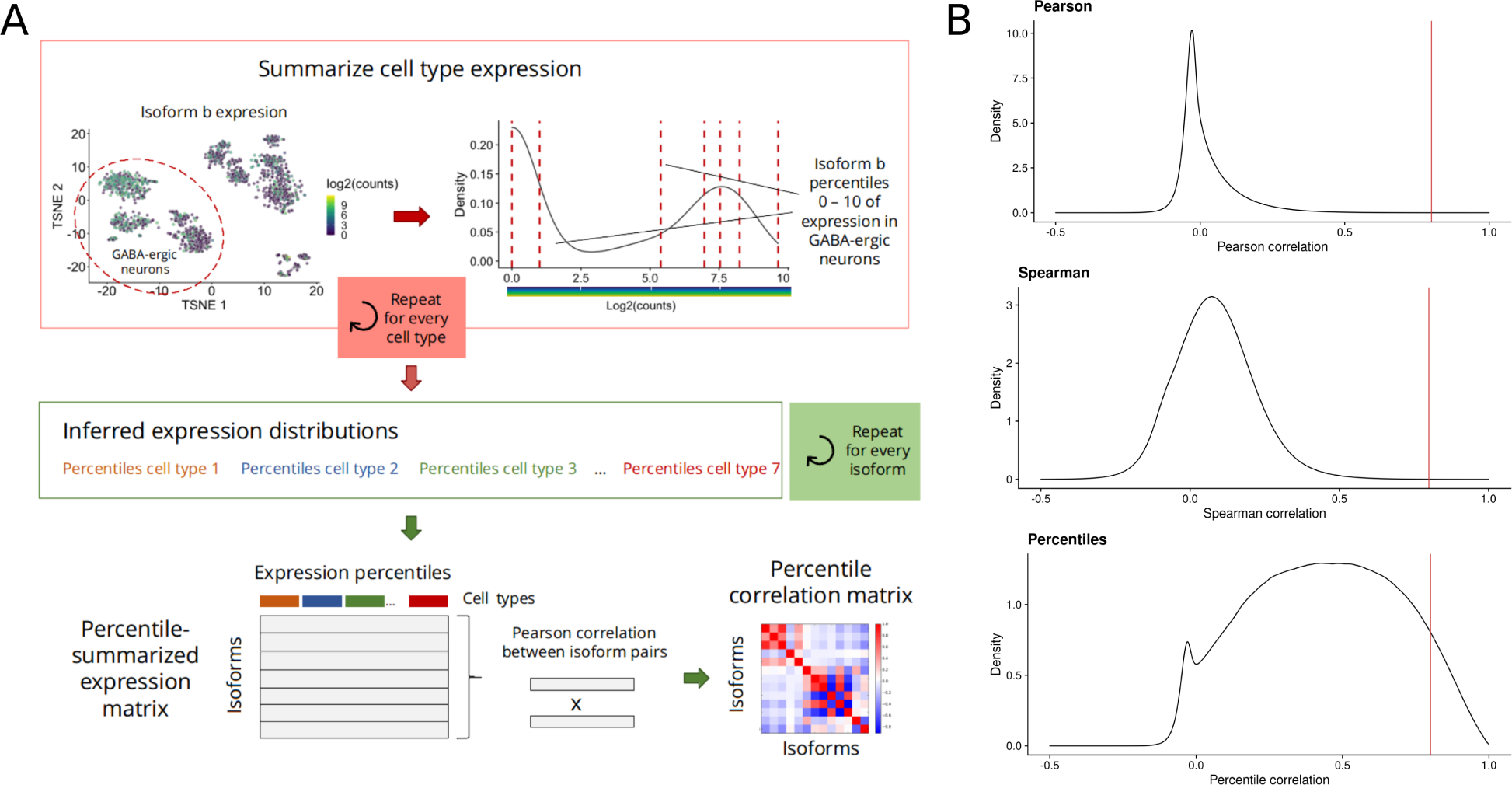
Percentile correlations. A) Percentile correlation algorithm. For each isoform, cell type-level expression is summarized using percentiles (0 - 10) as a proxy of the expression distribution of the isoform in each of the cell types. Then, Pearson correlations are computed using the percentile-summarized expression of all isoforms, obtaining a percentile correlation matrix. B) Correlation density distributions. Pairwise isoform correlations were computed using Pearson, Spearman and percentile+Pearson correlation. Red vertical line indicates the cor = 0.8 thres-hold.

Our approach considers cell-type identity to be defined by context-specific regulation of gene expression programs, and within-cell type stochasticity to arise from a combination of technical noise^52, 53^ and biological mechanisms such as transcriptional bursting^54^, which translate into the sparsity and heterogeneous expression patterns typically observed in scRNA-seq (Supplementary Figure 3A) and result in high variance across all levels of expression (Supplementary Figure 3B). Together, these effects mask the co-expression signal in the data and tend to yield low correlation values when using traditional metrics (Figure 2B). To overcome this problem, we treat single cells of the same cell-type as biological replicates that represent the state of a delimited cell population but are differently affected by the aforementioned combination of technical and biological forces. In this context, the expression distribution of any given transcript across the population can be considered as the signature of the transcript in that cell-type. To translate these assumptions into a metric, every isoform’s expression within a cell type is first summarized into an expression profile, where single-cell count values are replaced by 10 percentile values (deciles) (Figure 2A, Methods). Intuitively, this reduced number of values synthesizes the approximate behavior of that transcript in the cell type, as inferred given cell-level observations. Next, to grasp expression distribution similarities across cell types, Pearson correlation between transcript pairs is computed using the percentile-summarized expression, resulting in a meaningful distribution of correlation values (Figure 2B). In this manner, we extend the notion of cell-type markers to rely not only on mean or frequency of expression, but on their actual distributional pattern. Our co-expression metric therefore by-passes cell-level matching of individual observations, providing a correlation estimate that is both robust to the uncertainty of single cell expression and interpretable as a measure of expression similarity. Of note, changing percentile number did not have a noticeable effect on the resulting correlation values (Supplementary Figure 3C), which were considerably higher than those generated using traditional metrics, successfully unlocking co-expression analysis in single-cell data.

To detect modules of co-expressed isoforms, we used the percentile correlation matrix as a distance matrix for hierarchical clustering, and designed a semi-automated cluster refinement approach to ensure maximal profile similarity within clustered modules (Figure 3A, see Methods for a detailed description). Briefly, we first used the *dynamicTreeCut* R package^55^ to set adaptive thresholds and find clusters dynamically within the dendrogram. Among the resulting 152 clusters, some showed highly similar (i.e. redundant) expression profiles (Supplementary Figure 4A) and were therefore merged by re-clustering using the mean scaled expression of the isoforms in each cluster (i.e. clustering of metatranscripts, see Methods), obtaining 60 isoform clusters. Among them, 18 groups presented noisy expression profiles (Supplementary Figure 4B) and their isoforms were therefore re-assigned to the remaining 42 clusters. However, fully automated re-clustering was not effective to eliminate redundancy in some cases. As a result, conflicting cases were inspected and merged, generating a final number of 19 distinct clusters containing 8,688 isoforms (96,8% of total) and representing diverse expression modalities across the 7 broad cell types (Figure 3B).

**Figure 3:**
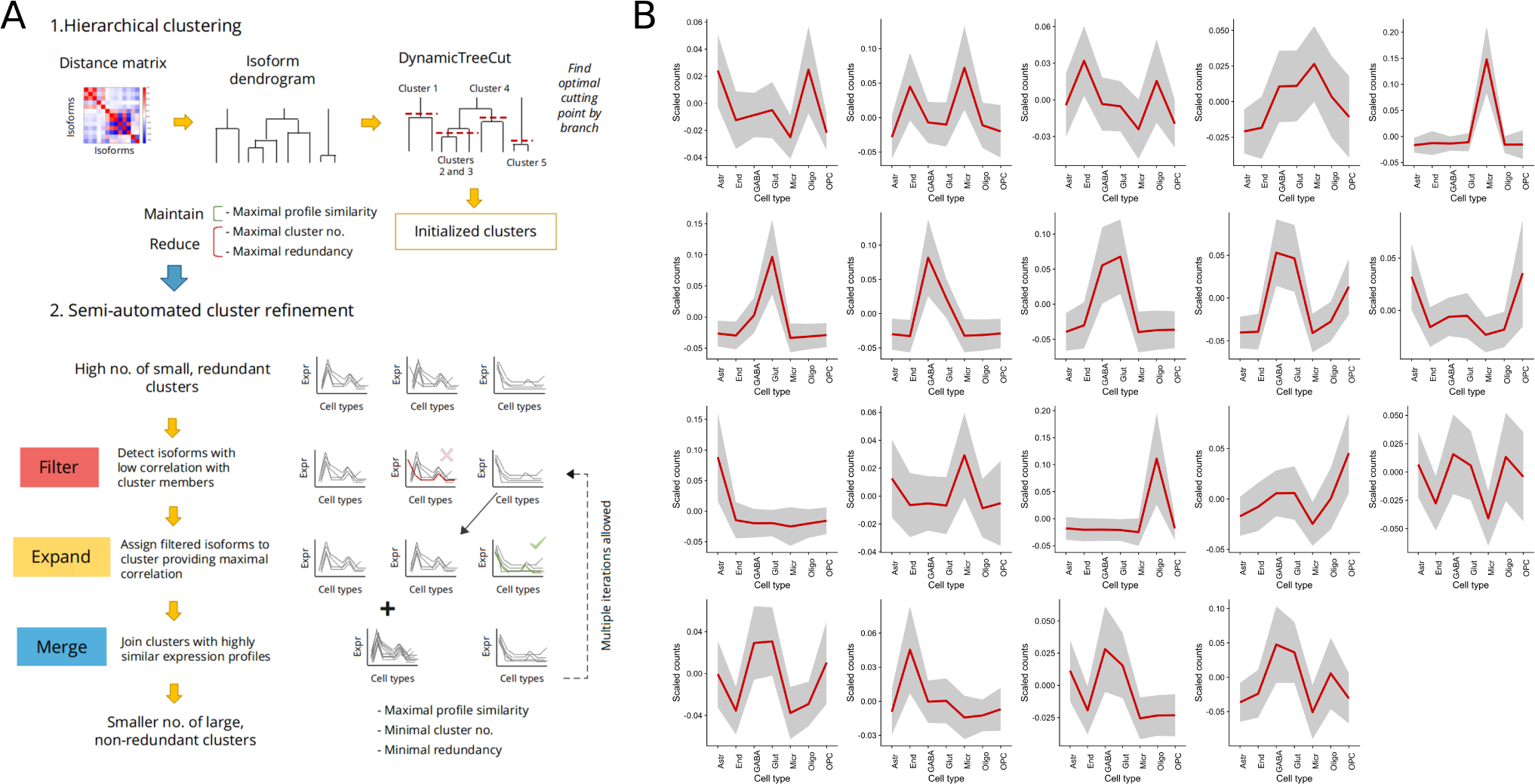
isoform clustering. A) Clustering pipeline. The percentile correlation matrix is first used as a distance matrix for hierarchical clustering. After dynamic cluster generation, noisy clusters are refined by a three-step semi-automated process. B) Clusters generated after applying the *acorde* clustering pipeline to the mouse neural data-set. Cell-level mean expression (scaled, see Methods) is computed for all transcripts and then aggregated as the global cell type mean, represented by the red line. Grey area corresponds to cell type mean ± standard deviation.

### Validation of percentile correlations on simulated data

Building on studies reporting the poor performance of popular correlation metrics in single-cell data, authors have attempted the implementation of sparsity-aware measurements^51, 56^ and reported the potential of other alternatives to compute similarity, such as proportionality metrics^42^. Here, we present an interpretable, scalable and biology-aware alternative to single-cell co-expression studies based on Pearson or Spearman correlation. However, to better understand how percentile correlation performs in comparison to extant correlation metrics, we compared it to Pearson, Spearman and zero-inflated Kendall correlations^56^ and one proportionality metric, rho (*ρ*)^57^ using simulated data.

Given that there are -to the best of our knowledge-no scRNA-seq data simulators that include transcript co-expression patterns, we designed a simulation strategy (Supplementary Figure 5A, Methods) to generate an appropriate validation framework for our metric. Briefly, we applied *SymSim*^58^ to simulate a single-cell RNA-seq dataset (8 cell types, 1,000 cells and 8,000 transcripts) and used the simulated expression values to artificially create 3,000 synthetic transcripts showing 15 different expression profiles across the 8 cell types (Supplementary Figure 5B). As a result, our simulated dataset contained 15 simulated clusters with distinct expression profiles. Among them, clusters 1-5, 6-10 and 11-15 included transcripts showing high expression in one, two and three cell types respectively, gradually increasing simulated pattern complexity (Supplementary Figures 5B-C). After refinement (Supplementary Figure 5C, Methods), 1,790 synthetic transcripts remained distributed across the 15 simulated clusters in groups ranging from 180 to 60 transcripts (Supplementary Figure 5D).

In order to evaluate how well the 5 co-expression methods recapitulated the simulated patterns, we computed these metrics for all synthetic transcript pairs in each simulated cluster (Supplementary Figure 5E). Among them, percentile correlation consistently yielded the best proportion of high within-cluster correlations followed by *ρ*, however, rather counter-intuitively, *ρ* had only an average performance when low-complexity patterns were provided, with less than 20% output proportionality values >0.8 within clusters 1-5. Shockingly, zero-inflated Kendall correlation, a single cell-tailored metric, failed to recapitulate the simulated co-expression profiles and showed a considerably lower proportion of high correlations within the simulated clusters than Pearson and Spearman correlations. As a result of this evaluation, we can confidently assume that percentile correlations are useful to detect co-variation patterns, yielding overall higher correlation values than all other considered metrics.

Of note, and similarly to real data, the simulated dataset showed no detectable effect when varying the number of percentiles used to compute percentile correlations (Supplementary Figure 6).

Next, we compared the ability of each co-expression metric to inform clustering and group synthetic transcripts with similar expression patterns. To achieve this, we run our clustering pipeline on the simulated isoforms using the 5 metrics as distance (Supplementary Figures 7-11). To enable benchmarking, clustering was automated to always generate a total number of 15 calculated clusters (see Methods). In order to evaluate which metric worked best to detect co-expressed transcript groups, we considered internal correlations between the transcripts in generated clusters. We observed that *ρ* and percentile-generated clusters, unlike the remaining co-expression metrics, presented consistently high levels of internal correlation (Figure 4A). Notably, the distribution of co-expression values obtained using percentiles was the most robust among the five metrics (Supplementary Figure 12). We next assessed how well the clusters generated using each co-expression metric (i.e. calculated clusters) recapitulated the simulated clusters. Calculated and simulated clusters were paired based on the similarities between their mean cluster profiles (Supplementary Figure 13, Methods), and the Jaccard index (JI) for each simulated-calculated pair was computed to measure the agreement in synthetic transcript assignment (Figure 4B). Interestingly, results were highly heterogeneous for most methods: even though a number of simulated co-expression groups were easily detected by most metrics, no method was able to fully recapitulate the simulated clusters, with *ρ* proportionality, Pearson and percentile correlations being the most accurate (Figure 4B). Zero-inflated Kendall and Spearman correlations, on the other hand, showed consistently low agreement with the simulated transcript groups. Finally, we considered the number of transcripts that remained unclustered (Figure 4C) before and after re-clustering unassigned transcripts (i.e. cluster expansion step, see Methods). Pearson correlation provided successful cluster assignment for practically all transcripts in the simulated dataset, especially when incorporating percentiles (Pearson: ∼10% unclustered before expansion, ∼1% after; percentile: ∼4% unclustered before expansion, 0% after), whilst the rest of metrics performed significantly worse, leaving 20-30% of transcripts unassigned even after cluster expansion, with proportionality (∼30% unclustered before expansion, ∼25% after) being the less optimal. Altogether, though *ρ* demonstrated good performance in many aspects of clustering, including intra-cluster correlation and agreement with the simulated clustering, it was outperformed by percentile correlation when globally considering all evaluated parameters (Figure 4D). In addition to the fact that *ρ* failed to control for unassigned transcripts, computing means and standard deviations of Jaccard indices across simulated-calculated pairs showed percentile and Pearson correlations as the most consistently accurate methods. All in all, our synthetic data evaluations showed that the percentile correlation approach performed well -and more consistently than *ρ* proportionality-in all the evaluated features, and visibly captured co-expression better than both traditional and zero inflation-aware correlation metrics.

**Figure 4:**
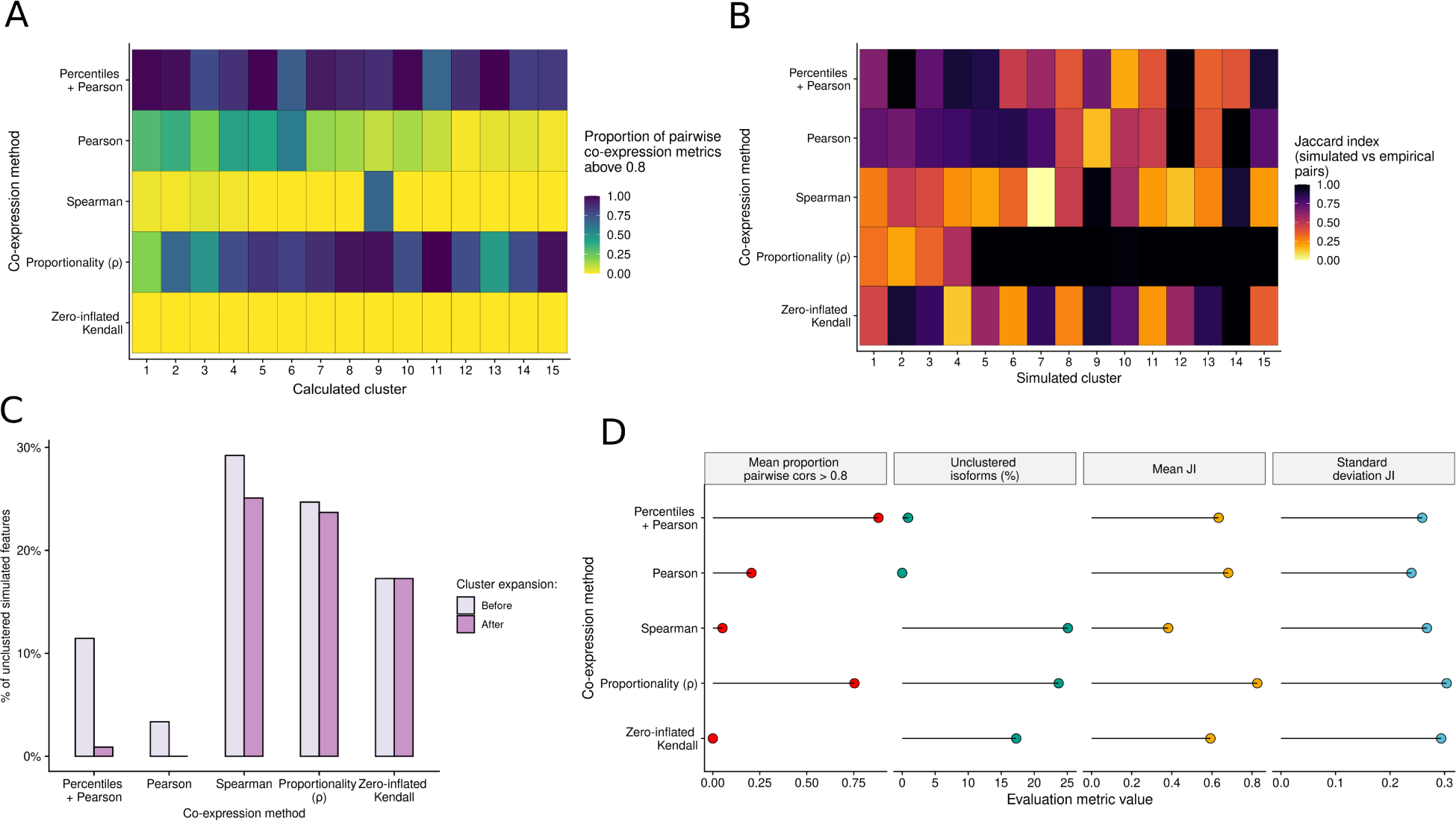
evaluation of percentile correlation-based clustering on simulated data. A) Proportion of co-expression values above 0.8 in each empirical cluster obtained after running the *acorde* clustering pipeline on simulated data using several correlation and proportionality methods as a distance metric. B) Jaccard Index of simulated vs calculated clusters obtained with each evaluated co-expression method. Simulated clusters were paired with one calculated cluster based on mean profile similarity, and synthetic transcripts present in each member of the pair were compared. C) Percentage of unclustered isoforms generated by each co-expression method, before and after re-assigning unclustered isoforms by measuring co-expression with the mean profile of extant clusters (i.e. cluster expansion). D) Evaluation metric overview. Metrics are specified in the grid headers. x-axis shows values of the different metrics, y-axis displays evaluated co-expression methods.

### Co-Differential Isoform Usage analysis of single-cell isoform expression

Isoform clusters represent groups of alternative transcripts that are co-expressed at the cell type level. However, our clustering results did not provide information on iso-transcriptome properties associated with splicing regulation. To facilitate interpretation of the isoform clustering results, we first defined genes with Differential Isoform Usage (DIU) as those whose isoforms were assigned to different clusters (Figure 5A) and found that 80% of genes with clustered isoforms (2,577 out of 3,172) were DIU and involved a total of 7,575 isoforms. In this context, DIU genes will necessarily have two or more isoforms with significant changes in expression across cell types and simultaneously undergo cell type-dependent post-transcriptional regulation, leading to changes in isoform expression in each cell type.

**Figure 5:**
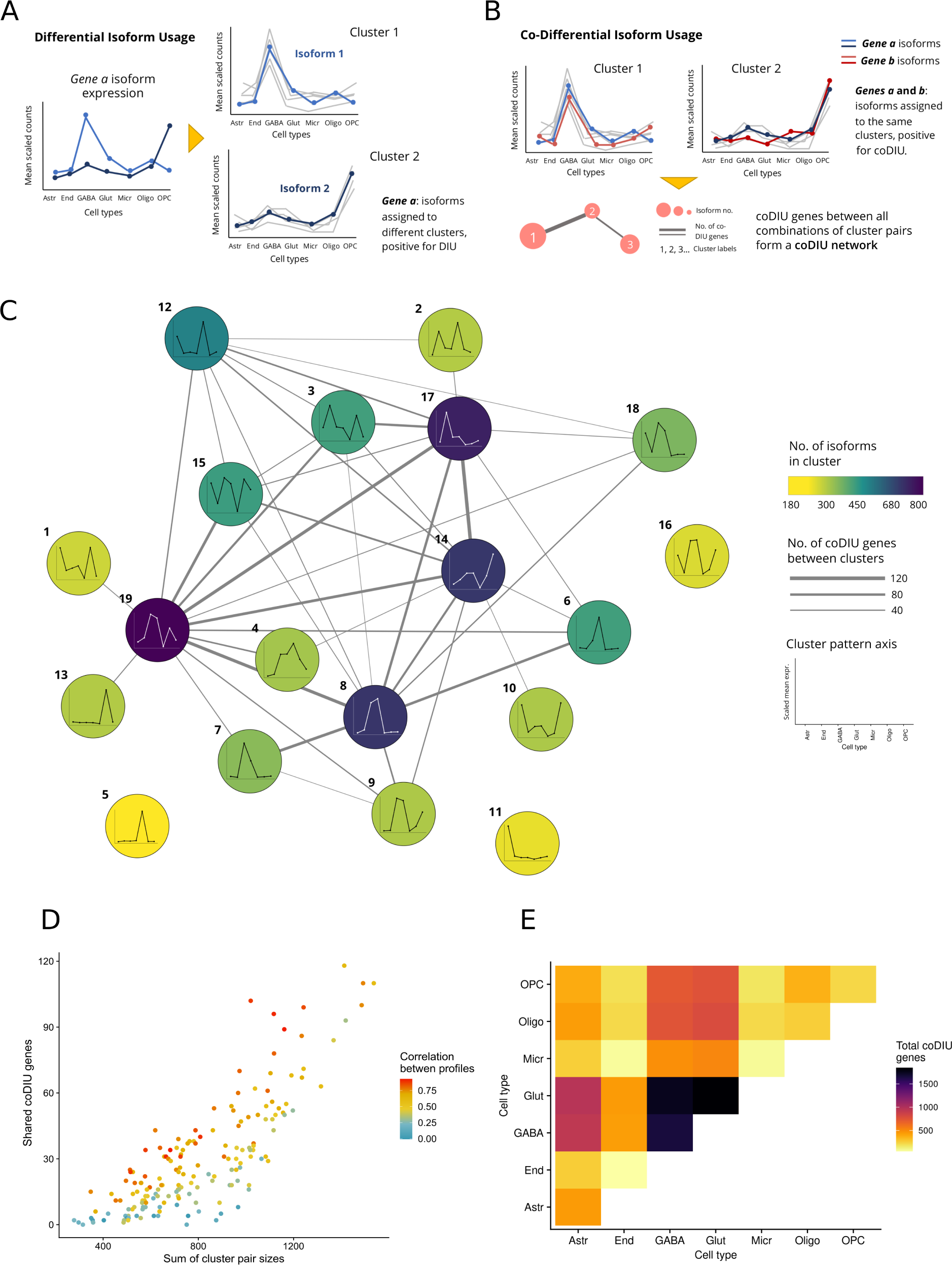
characterization of genes with co-Differential Isoform Usage. A) Cluster-based definition of Differential Isoform Usage (DIU) across multiple cell types. DIU genes that have at least two isoforms assigned to different clusters, indicating a differential isoform selection pattern across the different cell types. B) Definition of co-Differential Isoform Usage (coDIU) using clusters. CoDIU genes have multiple isoforms assigned to the same clusters, establishinig cross-cell type co-expression relationships for at least two of their isoforms. C) coDIU network. Nodes represent clusters and depict their mean expression profile across cell types. Node color represents cluster size (i.e. no. of isoforms in cluster). Edge width represents number of coDIU genes detected between each pair of clusters. D) Evaluation of cluster profile similarity and size as a function of the number of coDIU genes detected by *acorde*. x-axis corresponds to the sum of isoforms in each possible pair of clusters generated from the data. y-axis contains the number of coDIU genes between the pair. Dot color represents the correlation between the mean expression profiles of each pair of clusters. The number of coDIU genes between a pair of clusters is seemingly related to the size of the clusters involved, and shows no relationship with the degree of similarity betewen the expression profiles of clustered isoforms. E) Cell type-level coDIU patterns. For each pair of cell types represented in x and y-axis, heatmap color corresponds to the total number of genes found to be co-DIU between them. Total coDIU genes are calculated as the sum of coDIU genes detected between all occurrences of cluster pairs showing high expression of isoforms these cell types. GABA and Glut cell types share the highest number of coDIU genes, both with each other and with other cell types. Astr: astrcytes, End: endothelial cells, GABA: GABA-ergic neurons, Glut: glutamatergic neurons, Micr: microglia, Oligo: oligodendrocytes, OPC: oligodendrocyte precursor cells.

In order to study isoform co-expression patterns, we defined co-Differential Isoform Usage (coDIU) genes as those showing coordinated cell-type-specific isoform usage. Specifically, we considered two or more genes to be coDIU if their isoforms had been assigned to the same clusters (Figure 5B, Methods). This resulted in the definition of an isoform co-splicing network where nodes are clusters of correlated isoforms and edges represent coDIU genes, i.e. the number of genes for which two or more isoforms are co-expressed across cell types (Figure 5C, Methods). To ensure the selection of significantly coDIU genes, we fitted a generalized linear model for every pair of coDIU genes, and selected pairs with isoform-level co-expression and no significant cell type expression variation when only accounting for gene-level expression (see Methods). CoDIU genes therefore present cell type-dependent co-expression of at least two isoforms, represented by cluster assignment matches, but are not co-expressed when only gene expression is considered. Using this strategy, we detected 2,049 genes with at least one significant coDIU partner (*cluster*cell-type* FDR < 0.05, *gene*\**cell-type* FDR > 0.05), involving 6,370 co-expressed isoforms. The number of coDIU genes sharing isoforms across each cluster pair was variable, although it rose up to ∼120 for highly connected expression profiles (Figure 5C).

We then interrogated the coDIU network to find patterns underlying the splicing coordination signal detected by the *acorde* pipeline. First, in order to measure whether coDIU generated strong or subtle variations in isoform selection across cell types, we investigated the association of coDIU to single or multiple cell type isoform switching events. Note that single isoform switching events involve clusters with patterns that are similar across all cell types except one, leading to high between-cluster correlations. Interestingly, we found that the number of coDIU genes linking isoform co-expression clusters was dependent on cluster sizes, but showed no direct relationship with the similarities between expression profiles (Figure 5D). The detection of coDIU genes involving isoforms with highly different expression patterns suggested that coordinated isoform usage mechanisms are able produce strong cell type-level shifts in isoform selection, and thus modulate the expression of highly cell type-specific splice variants.

We next evaluated the cell type-level relationships present in the isoform co-expression network, namely the occurrence of coDIU across all possible pairs of cell types in our data. Although co-splicing could potentially occur between any combination of cell types, our results showed that a high proportion of coDIU interactions were detected when the isoforms involved had high expression in one of the two neural cell types, i.e. GABA-ergic and glutamatergic neurons (Figure 5E). This can be partially explained by the fact that some of the clusters with neuron expression are among the largest generated by the *acorde* pipeline (Figure 5C). However, another plausible explanation is that the central role of neurons in the tissue under study (i.e. primary visual cortex) might situate co-splicing at the core of neural function regulation, as well as the modulation of its interaction with glial cell types.

### Functional analysis of the coDIU network

We next set out to investigate the functional implications of our isoform co-expression network. Since, with a few exceptions^59, 60^, splicing analysis tools rarely integrate functional information and function annotation databases do not usually include data at isoform resolution, we annotated the long read-defined transcripts using IsoAnnotLite (https://isoannot.tappas.org/isoannot-lite/), included both transcript and protein-level motifs, sites and domains, as well as non-positional, gene-level features such as Gene Ontology (GO) terms (for a detailed description of the annotation process and a comprehensive list of functional categories and source databases, see Supplementary Note 1).

First, we analyzed which biological processes and gene functions were potentially controlled by DIU (AS-regulated) and coDIU (co-regulated) mechanisms operating across cell types, i.e. which functions were overrepresented at DIU and coDIU genes. In order to discriminate the functional properties of AS-regulated genes from those showing no cell type specificity in isoform expression, we performed a functional enrichment test of DIU genes vs genes with DE isoforms, which were used as the background (Figure 6A, see Methods). Interestingly, DIU genes showed significant enrichment (FDR < 0.05) of GO terms associated with transcription (*gene expression, DNA-templated transcription*) and RNA metabolism (*RNA processing, RNA metabolic process*), as well as protein features required for these processes, such as Nuclear Localization Signals (NLS), Post-Translational Modifications (PTMs), particularly acetylation sites (Acetyl-Lys), and protein-complex formation (Figure 6A). In order to investigate the cellular processes where coDIU could potentially have a relevant regulatory role, we compared the proportion of coDIU and DIU genes annotated for each functional feature in the transcriptome using a partially-overlapping samples z test^61^ (Figure 6B, see Methods). Not surprisingly, coDIU genes were significantly enriched (FDR<0.05) in some of the same functionalities obtained in the previous DIU analysis, such as RNA biosynthesis and processing (*transcription, gene expression, RNA metabolic process*). However -and in contrast to DIU genes, which were markedly involved in biosynthesis-, genes regulated by coDIU were erniched in mitochondria components (i.e. *mitochondrial matrix*), suggesting that coordinated isoform usage may affect oxidatoin and energy metabolism. Remarkably, coDIU genes also showed additional enrichment for splicing-related terms such as *RNA splicing*, *mRNA splicing via spliceosome* and for *3’ UTR motif K-box* (Figure 6B). This result links genes involved in splicing and elements affecting RNA stability with the coordination of AS and suggests that the co-expression of alternative isoforms is a post-transcriptionally regulated process, in line with previous evidence of AS self-regulation^62–65^.

**Figure 6:**
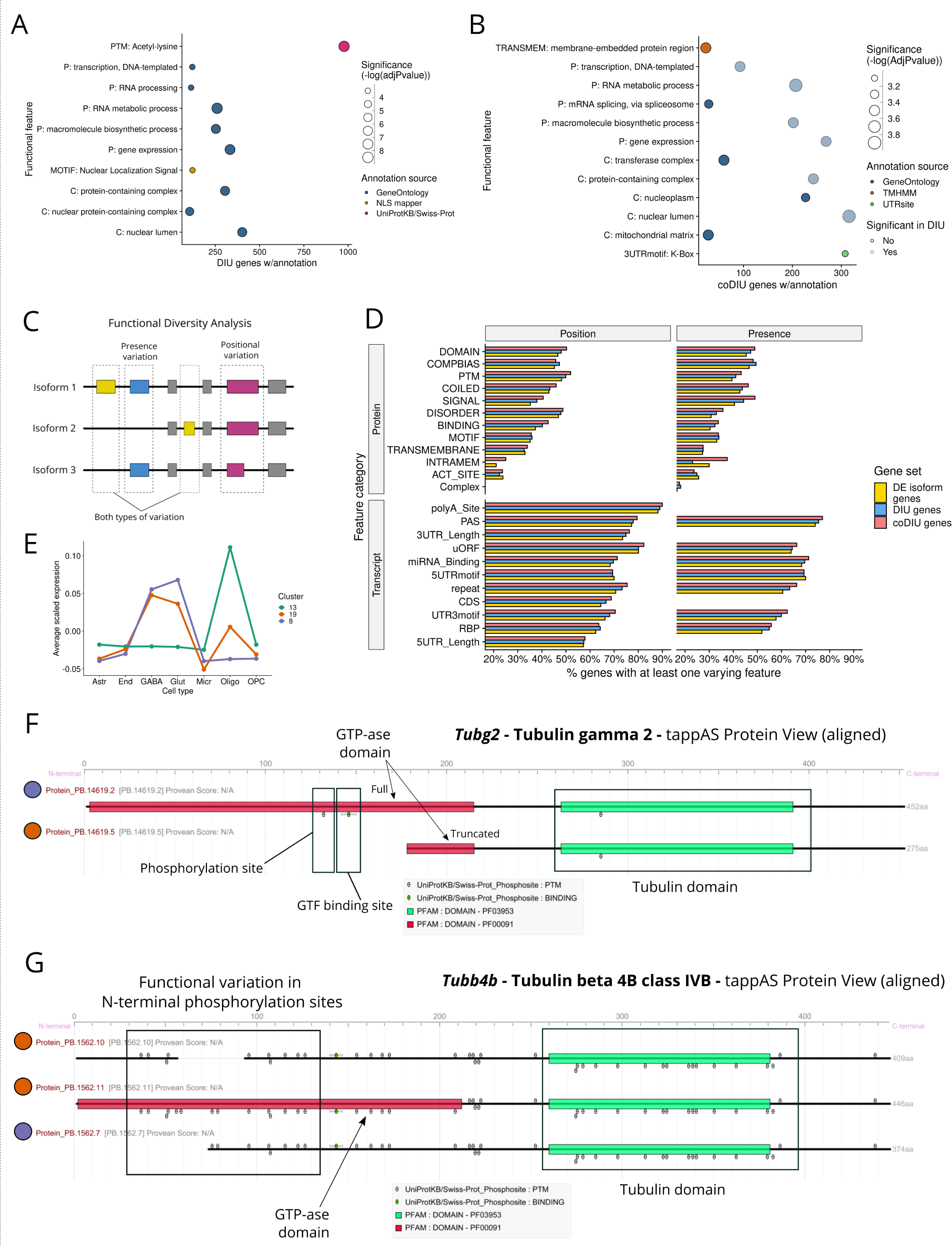
functional analysis of DIU and coDIU genes. Functional Enrichment results for A) Differential isoform Usage (DIU) genes vs genes with Differential Expression (DE) of isoforms and B) coDIU vs DIU genes. x-axis indicates the total number of test genes (A-DIU, B-coDIU) including the tested annotation feature, y-axis shows functional features. Dot color represents functional category, dot size represents -log(adjusted p-value). C) Schematic representation of the Functional Diversity Analysis (FDA). Variation using the genomic position and present/absence criteria are shown. D) FDA results for DE isoform, DIU and coDIU genes. y-axis shows transcript and protein functional categories (see Supplementary Note 1 for category definition information). x-axis shows the percentage of genes including at least one feature annotation from each of the categories that are detected as functionally *varying*. Both FDA criteria are shown (position - left grid column, presence - right grid column). CoDIU genes show the largest level of functional variation. E) Cell type expression patterns of clusters selected for downstream functional analysis: neural (cluster 8, purple), oligodendrocyte (cluster 13, green) or shared (cluster 19, orange). tappAS view of F) *Tub2g* and G) *Tubb4b* protein annotations. Cluster assignments for each isoform are indicated by dot color.

To obtain further insight into the functional elements controlled by AS co-regulation across neural cell types, we performed a Functional Diversity Analysis (FDA, Figure 6C). FDA is part of the tappAS framework^60^, and identifies functionally *varying* genes, i.e. genes expressing transcript variants with differences in the inclusion of functional features (see Methods). FDA can be evaluated from a presence/absence standpoint (i.e. AS completely removes a feature), or by detecting variation in the transcript positions defining the feature (Figure 6C). We therefore compared the diversity in transcript-level functional features between DIU, coDIU genes and genes with more than one DE isoform, for all functional categories provided by IsoAnnotLite (Figure 6D). Interestingly, we detected a larger percentage of varying genes as the level of considered regulatory complexity increased, with coDIU resulting in the largest amount of feature inclusion diversity in virtually all protein and transcript feature categories.

To measure this effect, we compared varying percentages between all combinations of the three gene sets (paired samples t-test, see Methods) and confirmed the observed trend, regardless of whether the variability criteria employed was position or presence. In particular, even though all comparisons were significant, coDIU resulted in the most significant increase in feature variation (coDIU vs DE isoform genes *p*-value: presence = 7.61e-07, position = 3.45e-05; coDIU vs DIU genes *p*-value: presence = 4.92e-04, position = 9.62e-03). We verified that this increase in functional diversity was not associated to expression level or isoform length biases in coDIU genes (Supplementary Figures 14A-B). This result suggests that alternative isoforms that engage in co-expression relationships tend to alter their functional properties significantly more often than other transcripts, namely by controlling the inclusion of motifs, sites and /or domains, thereby coupling alternative splicing and isoform co-expression with functional potential.

### Functional analysis of neuron-oligodendrocyte isoform co-expression

To further understand the relationship between cell-type identity, isoform co-expression and the functional properties of coDIU genes, we searched the coDIU network for cluster groups representing biologically-related isoform switches between defined neural types. Namely, we focused on a set of genes 160 coDIU genes (Figure 5C) representing oligodendrocyte-specific, neuron-specific, and shared isoform expression patterns (clusters 8, 13 and 19, respectively, Figure 6E) and analyzed isoform-associated functional variability using FDA. For this set of alternative isoforms, polyA site and 3’UTR length showed the highest variation rates among annotated transcript-level functional categories (varying in ≥70% genes, Supplementary Figure 15A). Moreover, we noticed that these changes followed a clear cell type-specific pattern, with the majority of coDIU genes shared among the three clusters expressing their longest 3’UTR isoforms in neurons (Supplementary Figure 15B) and neural-specific isoforms generally expressing longer 3’UTRs than their oligodendrocyte-specific counterparts (Supplementary Figure 15C).

Next, we inspected 3’UTR-related functional categories, i.e. RBP binding, miRNA binding, and 3’UTR motifs, using ID-level FDA (see Methods) to identify specific functional features associated to isoform usage differences between neurons and oligodendrocytes. Regarding the presence of RBP binding sites, we found that Mbnl and CELF4 binding sites showed high variation levels, i.e. 50% and 75% of genes, respectively (Supplementary Figure 15D); however, the amount of genes including Mbnl and CELF4 binding motifs was low (∼3% genes), possibly due to the relatively poor annotation density of RBP sites transferred by IsoAnnotLite (Supplementary Note 1). We also detected high variation frequencies for several miRNA sites (Supplementary Figure 15D), although no specific miRNA motif was shared by more than ∼5% of genes (up to 9 out of 160 genes). Nevertheless, regarding 3’UTR motifs, we found that the *K-box* motif presented inclusion changes in ∼60% of annotated coDIU genes (Figure 6A).

Interestingly, *K-box* motifs have been described to have a role in negative post-transcriptional regulation by promoting miRNA binding and transcript degradation^66, 67^. In line with this, the coDIU network included several genes in which 3’UTR elongation led to neuron-specific co-inclusion of *K-box* motifs, included *Atxn10* (ataxin 10, Supplementary Figure 16A) and *Btrc* (beta-transducin repeat containing gene, Supplementary Figure 16B), both of which have been proposed to be involved in neuron survival and differentiation^68, 69^. These results suggest that a 3’ UTR binding-mediated mechanism favoring isoform co-expression may operate to fine-tune the post-transcriptional regulation of neuron survival genes.

Importantly, the majority of neuron-oligodendrocyte coDIU genes also presented coding region variation (i.e. CDS, Supplementary Figure 15A), revealing that the coordination of isoform usage can modify both transcript and protein functional properties. In particular, protein domains (PFAM) and post-translational modifications (PTMs) presented high variation rates (varying in ∼40% of genes, Supplementary Figure 15A) and thus constituted the categories with the most cell type-level dependent functional variation. While ID-FDA reported no specific PFAM domains shared among the analyzed gene set (∼1%, maximum of 2 out of 160), up to ∼14% of them presented inclusion variation in similar PTMs (23 out of 160) with medium to high variation rates for phosphorylation, acetylation, ubiquitination and ligand binding sites (Supplementary Figure 15D). However, synergies between PTM and domain inclusion changes could still result in differential functional activities at the protein level. As an example, we found two tubulin isotypes, i.e. γ-tubulin 2 (*Tubg2*) and β-4-tubulin B-chain (*Tubb4b*), that co-expressed isoforms in clusters 8 and 19, that is, protein variants with neuron-specific and neuron-oligodendrocyte expression, respectively. Interestingly, both of these genes also presented inclusion changes in an N-terminal GTP-ase protein domain and several PTMs (Figure 6F and Figure 6G), although with different functional outcome. *Tubg2* presented neuron-specific expression of the full-domain, PTM-including isoform, resulting in the inclusion of a phosphoserine residue and a GTP binding motif (Figure 6F) and suggesting the GTP-ase role to be enhanced in mouse visual cortex neurons in comparison to oligodendrocytes. On the other hand, the domain-including *Tubb4b* isoform was expressed in both cell types (Figure 6G), suggesting that this β-tubulin isotype has a broader GTP-ase function than *Tubg2*. The domain/motif inclusion pattern arising from isoform co-expression in these two tubulin isotypes may reflect the different cellular roles of γ and *α/*β tubulins^70^. Specifically, *α/*β tubulins are the building-blocks of the microtubule cytoskeleton, while γ-tubulin has a major role in microtubule nucleation and on the regulation of microtubule dynamics. The γ-tubulins only present two isotypes, among which *Tubg2* is exclusive to neurons and has a well-defined role in neuron survival and growth^71^, whereas *Tubb4b* is part of a broader catalogue of β-tubulin isotypes, all of which are jointly regulated and share a structural function^72^. Intriguingly, however, mutations in a closely-related isotype (*Tubb4a*) have already been shown to have a differential phenotypic effect in neurons and oligodendrocytes^73^. In line with this, we observed modifications in the number and density of N-terminal phosphorylation sites in *Tubb4b* as a result of cell type-level isoform co-expression, with an increase in protein diversity in neurons (3 vs 2 alternative isoforms, Figure 16G). Even more interestingly, PTM changes on different *α*/β-tubulin isotypes have long been known to modify tubulin stability and its interactions with other proteins^74^, creating a “tubulin code”^75^.

As a result, tubulin PTMs have the ability to modulate microtubule stability and function, intervening in processes such as neural differentiation, survival and polarity^76, 77^. This cell type-specific inclusion of different PTMs via isoform co-expression could therefore operate as a fine-tuning mechanism of the tubulin code, modifying the number and position of available modification sites in β-4-tubulin.

## Discussion

Alternative Splicing (AS) is known to be a tightly-regulated process in which splicing factors interact to create cell type-specific isoform expression patterns^78^. The transcriptome-level consequences of AS regulation have been studied in different ways, including, but not limited to, the detection of within-isoform coordination of alternative sites^40, 41^, the generation of gene-isoform networks to uncover novel regulatory relationships^79–82^ and the application of single-cell data to unravel cell type-specific expression patterns for same-gene isoforms^27, 83^. However, the extent to which AS regulation creates co-expression patterns among alternative isoforms from different genes has not yet been fully addressed. Specifically, previous studies tackling this type of isoform co-expression have either focused on specific event types, such as alternative 3’ exons^84^, or solely on the identification of functionally-relevant alternative isoforms in different biological contexts^85, 86^.

In this study we present *acorde*, an end-to-end pipeline to generate isoform co-expression networks and detect genes with co-Differential Isoform Usage (coDIU), and apply it to the study of isoform co-expression among seven neural broad cell types^43^. To this end, we successfully leveraged single-cell data by implementing percentile correlations, a metric designed to overcome single-cell noise and sparsity and provide high-confidence estimates of isoform-to-isoform correlation. Here, we show that percentile-summarized Pearson correlations outperform both classic and single-cell specific correlation strategies^56^, including proportionality methods that were recently proposed as one of the best alternatives to measure co-expression in single-cell data^42^. In addition, using a long read-defined, functionally annotated transcriptome enabled us to obtain a biological readout from the isoform network. coDIU genes were found to be enriched in the same biological functions, a number of which were unique in comparison to enrichment results for genes solely reported as DIU. Inter-gene isoform co-expression thus appears to impact a subset of DIU genes sharing a specific set functions, which suggests that coDIU may contribute with an additional layer of complexity to some cellular processes, operating as a fine-tuning mechanism. Our analyses also revealed that isoforms from coDIU genes encompass higher functional diversity than those belonging to DIU genes, an effect that was not associated to expression or length differences.

Importantly, these changes impact both transcript and protein isoform functional features, which pinpoints their ability to globally increase the functional repertoire of coDIU genes. Mechanisms generating isoform co-expression can therefore be thought of as a potential source of functional synergies between alternatively spliced genes, giving rise to simultaneous changes in functional properties among co-expressed isoforms.

To demonstrate the power of the *acorde* pipeline, we include examples where these kinds of coordinated changes were detected in the mouse neural dataset. First, we report a neural-specific pattern of 3’UTR co-elongation that is consistent with the available literature^87, 88^ and results in a simultaneous increase in the number of K-box motifs, miRNA and RBP binding sites in these UTRs. In addition, we describe how cell type-specific co-expression of γ and β-tubulin isoforms creates divergent functional properties among them, which is consistent with their different biological functions^71, 72^ and points to a splicing-mediated coordination of cell-type specific structural components. On a broader note, an interesting finding stemming from our functional analyses concerns the type of biological properties that make the functional signal of the coDIU network. Namely, while we do find enrichment of biological processes (i.e. *RNA metabolism* or *mRNA splicing via spliceosome*) and differential inclusion of many features from a wide variety of functional categories, we failed to recover specific functional elements consistently present in a large number of co-expressed isoforms, such as binding sites for specific RBPs or microRNAs. This suggests that AS and post-transcriptional regulation mechanisms operate by controlling the presence of a broad array of functional features, which jointly contribute to modulate key cell-level processes encoding cell-type identity. While these insights need to be subject to further experimental validation, they serve to illustrate the hypothesis-generating power of our pipeline.

All in all, we have hereby showed that *acorde* can effectively leverage single-cell RNA-seq data to build isoform co-expression networks, revealing a new dimension of post-transcriptional regulation of gene expression while also disclosing the cellular processes and functional elements impacted by these mechanisms.

## Methods

### Single-cell data pre-processing and quality control

Mouse neural single-cell RNA-Seq data from mouse primary visual cortex was obtained from Tasic *et al.*^43^ and consists in paired-end Illumina reads generated with the Smart-seq2 protocol^89^, which enables isoform-level quantification. Reads were downloaded from Sequence Read Archive accession SRP061902 and mapped to the mouse genome (GRCm38.p6) using STAR^90^. We performed isoform expression quantification of the long read-defined isoforms (see Supplementary Note 1 for details on long read transcriptome definition) using RSEM^91^, and used the labels provided by Tasic *et al.* to assign the 1679 cells to 7 broad cell types: 5 glial (microglia, endothelial cells, oligodendrocytes, oligodendrocyte precursor cells (OPCs) and astrocytes) and 2 neural (GABA-ergic and glutamatergic neurons).

Isoform length effect on expression was evaluated using the *NOISeq* R package^92^, where mean expression showed to be highly correlated with transcript length (adjusted R^2^ = 0.81; p-value = 2.2e-16). Using isoform *i* effective length (*l_i_*) and cell-level *j* estimated counts (*c_ij_*), both output by RSEM and, after testing several alternatives, we devised a custom formula to minimize the impact of length on isoform expression for each isoform *i*:

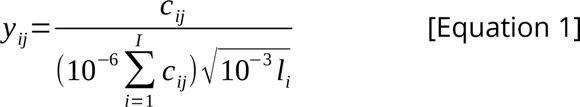

The transformed expression value for isoform *i* in cell *j* (*y_ij_*) was again tested for length bias and a low correlation was found (adjusted R^2^ = 0.25; p-value = 6.84e-8). Next, we inspected the library size distribution and filtered both high and low-count outliers due to potential premature cell death or library preparation duplets, with a total of 1591 cells passing quality control. Feature-level quality control was performed in a cell type-aware manner, keeping isoforms that showed non-zero expression in at least 20% of one cell type. Out of the 36,986 isoforms and 12,692 genes in the PacBio-defined transcriptome, we retained 18,867 isoforms and 9,596 genes for downstream analysis. Finally, we retained isoforms from multiple-isoform genes, hence removing cases where no AS-directed, isoform-level co-expression relationships can be established. This sets a general requirement for the entire study, which is that all isoforms retained must have, at all times, at least one same-gene counterpart to establish regulatory relationships that can be based on differential splicing of that gene, given that no AS regulation can be detected if a gene’s total expression is represented by a single isoform.

As a result, 13,832 isoforms from 4,591 genes were preserved.

### Differential Expression (DE) across multiple groups

#### Single-cell DE analysis

Differential Expression (DE) analysis among the 7 cell types was performed by combining ZinBWaVE weights^47^ and bulk-designed DE methods edgeR^49^ and DESeq2^48^ (i.e. using the corresponding R packages), which enable multiple group testing and were among the best-performing methods when combined with the ZinBWaVE method. Briefly, ZinBWaVE calculates cell-level weights for each isoform, effectively downweighting zeros during modelling for differential expression in single cell data (see van den Berge *et al.*^47^ for details), and hence unlocking bulk RNA-Seq computational methods for single-cell data. Of note, generalized linear models (GLM) within edgeR and DESeq2 were built and run following the pipeline used by van den Berge et al.^47^ to make them suitable for single-cell RNA-seq data, and are implemented in a wrapper function within our R package as described in Equation 2, where *y_ij_* is expression of isoform *i* in cell *j*, *T_kj_* is a dummy variable which takes value 1 when cell *j* is assigned to cell type *k* (*k*=1,…,*K*) and 0 otherwise, *βki* are the regression coefficients for isoform *i*, *εij* represents the error term, and *h()* is the link function of the GLM (natural logarithm in this case).

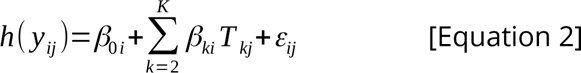

Differential Expression was defined using a significance threshold of FDR<0.05 when testing the significance of the model for each isoform *i*, that is, *H0: β2i = … = βKi.* Isoforms considered DE were preserved for downstream analysis if detected by at least one of the two methods edgeR or DESeq2, since this indicates a change in expression for any of the cell types considered rather than a flat expression profile. In addition, isoforms were removed if they belonged to genes with a single DE isoform, in agreement with the idea that no differential splicing regulation can be detected for single isoform genes, and hence that they will not form co-splicing relationships with isoforms from other genes in the network.

Prior to DE testing, and to balance the number of cells per cell type, the most abundant cell types (GABA-ergic neurons, n = 761; glutamatergic neurons, n = 764 cells) were downsampled by randomly selecting 45 cells, keeping N=241 cells for downstream analysis, simultaneously reducing the computational cost of multi-group DE testing. Finally, since downsampling can lead to having isoforms with all-zero counts again, we re-filtered isoforms that retained non-zero expression in at least 20% of one cell type, ultimately keeping 13,451 isoforms from 4,583 genes for the DE analysis.

#### Validation of downsampling and single-cell DE analysis

To check that the random selection of cells in the downsampling process was not affecting the DE results, we performed 50 independent runs of random neural cell sampling (without replacement) followed by zero-expression filtering (expression above zero for at least 25% of cells in at least one cell type, where threshold stringency was increased with respect to the standard pipeline to best control zero abundance among iterations and avoid problems during GLM modelling) and DE testing with edgeR and DESeq2. First, to measure the consistency of each method independently, we calculated the mean and standard deviation of the number of DE isoforms across all same-method sampling runs (*R* = 50). To check the level of within-method agreement, we next considered isoform IDs labeled as DE in each independent method run, and calculated the Jaccard Index (*J_rs_*) between DE results of that same method for all possible pairs of random sampling runs *r* and s (*r*,*s* =1,…,50, *r*<*s*, a total of 1225 comparisons). To summarize this information, we relied on the mean and standard deviation of these two sets of *J_rs_* values. Finally, we measured the level of agreement between edgeR and DESeq2 regarding our DE criteria, that is, considering isoforms detected by at least one of the methods to be significantly DE (FDR<0.05). To achieve this, we calculated the union of DE isoforms between one-to-one pairs of edgeR and DESeq2 runs (*R* = 50), and computed the Jaccard Index between all possible pairwise combination of global DE results, i.e. isoforms detected by at least one method (again, 1,225 comparisons).

### Percentile correlation

In order to assess the similarity of isoform expression profiles across cells, a correlation measurement can be used by taking cells as observations. We propose here instead to first summarize the expression within a given cell type with percentiles and then compute the correlation using all cell types and their percentiles as observations, a method that we refer to as “percentile correlation” (see main text Figure 2A).

Percentile correlations rely on the assumption that cell-to-cell differences can be mostly attributed to transcriptional stochasticity or technical noise, and that these within-cell type differences have a smaller effect than between-cell type expression differences. However, expression estimates for transcripts within the same cell are biased in different degrees, mostly depending on their expression levels, with lower expression being generally accompanied by higher noise levels^52^. This modifies the extent to which isoforms are affected by noise in each cell and causes strong cell-level effects that prevent the detection of co-expression relationships using solely cell-level measurements. Instead, we set out to target changes in expression across cell groups. We therefore considered isoform expression levels in the different cell types as a range of possible values, defined by the cell-level measurements in the data. In this context, the expression value of an isoform in a cell is used a proxy to infer the underlying distribution of expression values in the cell type, where the shape and width of this distribution will depend on both biological and technical factors.

To translate this into a metric, we first took the expression values of an isoform in each of the cell types and computed a number of percentiles (*p*). We selected *p =* 10 to achieve a good balance between accuracy and computational burden in downstream analysis. As the minimum expression value (percentile 0) was also included, we actually had 11 new values representing the expression range within a given cell type. As a result, each isoform will possess a new, recalculated expression vector where the percentile values computed in each cell type will replace cell-level expression estimates. This process was repeated for each isoform. Next, we computed pairwise Pearson correlations between every pair of isoforms, obtaining a percentile correlation matrix **R**. In this context, high correlations will appear if a pair of isoforms shows a similarly broad expression distribution in most cell types, as well as a similar amount of relative expression change between cell types.

### Semi-automated isoform clustering

In order to obtain modules of tightly co-expressed isoforms, we combined the hierarchical clustering algorithm with several rounds of cluster profile refinement (see main text, Figure 3A), in order to automate the most intensive steps of clustering while also granting control over the level of aggregation and within-cluster similarity. Clustering and refinement steps can be combined and re-arranged to best capture co-expression patterns within the data, and their parameters can be defined by potential future users to provide maximum flexibility.

Functions for clustering and refinement are implemented in the *acorde* R package.

#### Dynamic hierarchical clustering

The previously obtained correlation matrix (**R**), where each element rij represents the Pearson’s correlation coefficient between the percentiles of isoforms (*i*,*j*), was transformed into a distance metric to be used in the hierarchical clustering. As we aimed to cluster positively correlated isoforms given our biological hypothesis, we discarded negative correlation values by replacing them with zero values, and therefore defined the distance between any pair of isoforms *i* and *j* as in Equation 3.

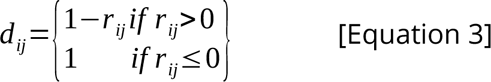

We next performed hierarchical cluster analysis using the *hclust()* function in the R *base* package^93^ with the average linkage criterion, and obtained a dendrogram. To obtain clusters, we used the *cutreeHybrid()* function in the *dynamicTreeCut* R package^55^ in order to find different thresholds for different branches of the dendrogram tree, instead of using a fixed threshold for the entire dendrogram. We provided the following non-default parameters to the *cutreeHybrid()* function: *deepSplit = 4, pamStage = FALSE, minClusterSize = 20*. Briefly, *deepSplit* ranges between 0 and 4, and provides smaller clusters, more accurate clusters when set to high values. *pamStage* determines whether a second stage of clustering using an algorithm similar to the Partition Around Medoids (PAM) method will be performed after searching the dendrogram for clusters (see Langfelder *et al*.^55^). As a result of this PAM-like step, no items are left unassigned to clusters, while setting pamStage to FALSE allows unclustered items. Finally, *minClusterSize* determines the minimum size of the produced clusters, and thus passing a higher value to this argument prevents the generation of too many clusters with a very small number of items.

This initial set of clusters is to be used as “hooks” to gather as much expression profile diversity from the data as possible. Importantly, even though our parametrization allows isoforms to remain unassigned to clusters (see above), some isoforms may still show low similarity to their cluster’s profile. To be able to obtain profiles as consistent as possible for downstream refinement, we implemented a cluster quality control step in which we removed isoforms based on a minimum correlation threshold with the rest of the members. In particular, isoforms were moved to the unclustered group if they showed a correlation lower than 0.85 with 3 or more isoforms from their cluster. In this manner, only tightly-correlated groups of isoforms will remain clustered.

#### Expanding clusters with unassigned isoforms

To re-assign unclustered isoforms to clusters with which they show high correlation, we performed two rounds of correlation-based, cluster expansion. In this process, we first summarized each cluster’s profile into a synthetic representative transcript that we named “metatranscript”. Metatranscripts were calculated as the mean of the percentile-based expression of all isoforms in the cluster. As a result, we obtained 11·*K* (*K* being the number of cell types) mean-summarized percentile expression values, which can be understood as an approximation to the expression range shown by the isoforms from that cluster in each of the cell types.

We next computed correlations between metatranscripts and unclustered isoforms, and performed two rounds of assignment. First, we assigned unclustered isoforms showing correlation values above 0.9 with at least one cluster, where the maximally correlated cluster was selected as the best match if there were ties. Next, metatranscripts were re-calculated for the newly expanded clusters and a similar assignment round was performed, this time lowering the correlation threshold to 0.8. In doing this, unclustered isoform groups are assigned in order, and highly correlated elements can therefore contribute to strengthen within-cluster similarities before assigning more lowly correlated elements.

#### Merging clusters by profile similarity

Prioritizing the reduction of within-cluster variability led to obtaining a large number of small, redundant clusters. To mitigate this effect while also preserving high correlations between cluster members, we merged clusters by profile similarity using the correlations between their metatranscripts. To achieve this, we performed dynamic hierarchical clustering using the *cutreeHybrid()* function on metatranscripts, with the following non-default parameters: *deepSplit = 4, pamStage = FALSE* (see section on dynamic hierarchical clustering above for more information), *minClusterSize = 1*. Of note, *minClusterSize* is set to 1 to avoid forcing merge of clusters that do not show enough similarity. As a result, we joined highly similar clusters, decreasing the number of clusters and hence minimizing redundancy.

While this strategy effectively eliminated between-cluster redundancy, we observed some exceptions where clearly similar profiles were not merged by clustering metatranscripts (Supplementary Figure 3A). In addition, a few cases arose where automated merging resulted in joining clusters with highly uncorrelated profiles (Supplementary Figure 3B). Given that it is very hard to provide a completely automated solution to the clustering problem, we solved these final discrepancies using a series of manually supervised refinement steps.

For that, we first inspected cluster profiles to flag incorrect merge decisions generating noisy clusters. Isoforms in these clusters were treated as unclustered, and used to expand the remaining, well-defined clusters. To do this, we computed the correlation between the isoforms to be assigned and cluster metatranscripts, similarly to the cluster expansion process described above. In this case, we joined isoforms to clusters if they presented Pearson correlation > 0.8 with its metatranscript. Isoforms showing high correlation with several clusters were assigned to the cluster with maximum correlation. Once isoforms from noisy clusters had been re-assigned, clusters were manually inspected again to detect profile redundancies missed by automated metatranscript clustering. Clusters that presented a similar expression pattern across cell types were then merged.

### Co-expression pattern simulation

To validate percentile correlations and our clustering strategy, we evaluated their performance on synthetic data, where co-expression relationships between simulated features need to be pre-defined as part of the data simulation process. However, there is, to the best of our knowledge, no currently available strategy to simulate single-cell data including modules of co-expressed features. We therefore designed our own simulation strategy by combining the *SymSim* R package^58^ to adequately model single-cell RNA-Seq data, and a dedicated strategy to generate co-expression between *SymSim* simulated features.

First, we set the following parameters to the *SimulateTrueCounts()* function in *SymSim* in order to obtain a count matrix consisting in 1000 cells from 8 cell types and 8000 features, with sufficient feature-level variation between the different cell groups:

SimulateTrueCounts(ncells_total = 1000, min_popsize = 100, i_minpop = 1, ngenes = 8000, nevf = 10, n_de_evf = 9, evf_type = “discrete”, phyla = pbtree(n = 7, type = “discrete), vary = “s”, Sigma = 0.25, gene_effect_prob = 0.5, bimod = 0.4, prop_hge = 0.03, mean_hge = 5)

Next, we modeled technical effects on these true counts in order to obtain real, observed counts using the *True2ObservedCounts()* function in *SymSim*, with the following parameters:

True2ObservedCounts(true_counts$counts, meta_cell = true_counts$cell_meta, protocol = “nonUMI”, alpha_mean = 0.1, alpha_sd = 0.005, lenslope = 0, gene_len = rep(1000, nrow(true_counts$counts)), depth_mean = 4e6, depth_sd = 1e4)

To create co-expression patterns, we then re-ranked expression values on a cell type-specific manner to define synthetic features, based on the expression profile of 15 pre-defined co-expression modules.

First, we drafted 15 different co-expression profiles reflecting three levels of expression complexity, that is, showing high expression or expression “peaks” in one, two, or three cell groups, respectively. To generate a count matrix reflecting these expression patterns, we shuffled simulated counts to create new, synthetic features. To achieve this, we first re-ranked features in each cell group by mean expression across cells in the group, breaking feature connectivity between the simulated cell types. Then, the top 1,400 features from each cell type were selected, together with the bottom 1,400 features. In this manner, we obtained high-expression and low-expression count vectors for each group, which we then combined to create synthetic features following the pre-designed cluster’s co-expression pattern. For each cluster, 200 count vectors from top-expression features were assigned to peaking groups, and 200 count vectors form bottom-expression features to cell groups showing low expression. Of note, 1,400 features were selected for simulation in order to grant at least 7 different 200-feature groups could be generated for each cell type, where the cell group with the highest peaking frequency across clusters showed high expression in 6 clusters only.

All in all, we obtained a simulated count matrix containing 1,000 cells from 8 cell types and 3,000 synthetic features, all of which belong to one of the 15 simulated co-expression modules. Therefore, by breaking feature-level connectivity between cell types, we benefited from feature-specific properties at the cell type level, while re-creating cell type expression coordination patterns that the SymSim strategy was not able to generate. Finally, to ensure the quality of the simulated clusters, we filtered synthetic features if their Pearson correlation with the cluster’s median profile was below 0.75 (Supplementary Figure 4B).

### Benchmarking of isoform correlation metrics for scRNA-seq data

Traditional correlation metrics have been shown to perform poorly when applied to scRNA-seq data, mainly given the increased noise and stochasticity levels in this data type. Recently, extensive benchmarks including single cell-tailored metrics have shed light on how to best select correlation metrics for single-cell data (see review by Skinnider *et al.*^42^). We therefore compared the performance of percentile correlations to a representative set of correlation metrics used in single-cell co-expression studies, namely classical Pearson and Spearman correlations, single-cell designed zero-inflated Kendall^56^ correlation, and proportionality metric rho (*ρ*)^57^, in agreement with previous reports showing that proportionality metrics were among the best performing co-expression methods in single-cell data. To measure performance, we computed these five co-expression metrics for all the synthetic features in the previously-simulated dataset, generating five different distance matrices for clustering, and evaluated which metric best recapitulated the simulated co-expression modules when used in our clustering pipeline. Pearson and Spearman correlations were computed using the *cor()* function in the R-base *stats* package. Zero-inflated Kendall correlation and rho (*ρ*) were computed using the *dismay()* function in the *dismay* R package^42^.

To make our benchmarking comparable, we adapted our clustering pipeline to remove all non-automated steps and always generate a fixed number of clusters. First, hierarchical clustering was performed on each correlation matrix using *dynamicTreeCut()*^55^ and the following non-default parameters to maximize granularity: *deepSplit = 4, pamStage = FALSE, minClusterSize = 10*. Of note, we skipped the quality filtering step based on intra-cluster correlations (see isoform clustering section above) to avoid bias against metrics that tend to yield low values when applied to single-cell data. Since we intended to evaluate the number of features remaining unclustered using each metric, we additionally suppressed the unclustered isoform assignment step (see isoform clustering section above). Finally, the merge process was automated by using the traditional hierarchical clustering algorithm (implemented in the *hclust()* function in the R base package) to group clusters based on the inferred metatranscripts that summarize the cluster’s expression profile (see isoform clustering section above). Finally, we set the number of clusters to 15, i.e. the number of simulated co-expression modules.

In addition to the number of unclustered isoforms, we used the levels of internal correlation in the empirical clusters, i.e. those obtained by *de novo* clustering of simulated synthetic features, to evaluate the clustering. We did this by aggregately considering all pairwise metric values for features within a cluster and measuring the percentage of metrics that are above a threshold value of 0.8. To assess how well empirical clusters recapitulated the co-expression simulation, we paired empirical with simulated clusters using the correlations between their mean cluster profiles. Simulated clusters were therefore paired with the empirical clustering showing maximum profile correlation. We next compared synthetic feature IDs assigned to the obtained and empirical clusters in each pair using the Jaccard Index (JI).

### Differential Isoform Usage and co-Differential Isoform Usage across multiple groups

#### Defining Differential Isoform Usage across multiple groups

Grouping isoforms into different clusters allows detection of a number of expression patterns across the multiple cell types included in single-cell data. As previously described, we filtered DE isoforms to ensure that all transcripts had at least one other counterpart from the same gene that was also significantly DE. Intuitively, in order for Differential Isoform Usage (DIU) to occur, a gene must first have at least two DE isoforms. However, we only considered a gene to be positive for DIU if (at least) two isoforms were DE and were assigned to different clusters, indicating that two of the gene’s isoforms show different expression patterns across groups (see main text, Figure 5A). Ultimately, this can be interpreted as an indicator that isoform expression regulation is cell type-dependent in that gene.

#### Detecting co-splicing patterns across isoform clusters: co-Differential Isoform Usage

We define coordinated splicing patterns as a situation where post-transcriptional regulation, defined by isoform expression, can be detected independently of transcriptional regulation, i.e. gene-level expression. To detect splicing coordination, we defined co-Differential Isoform Usage (coDIU) as a pattern where a group of genes shows co-expression of their isoforms, but no co-expression can be detected when only gene expression is considered (see main text, Figure 5B). In the context of our pipeline, a set of potentially coDIU genes will have at least two of their isoforms assigned to the same clusters, therefore showing detectable isoform-level co-expression, and suggesting coordinated splicing regulation in that group of genes. However, clustering allows expression pattern variability among members, and therefore some isoforms might be assigned to clusters that do not faithfully represent their expression profile, leading to detection of false-positive coDIU genes.

To identify groups of genes candidates for coDIU, we applied negative-binomial generalized linear regression models. Let *G* be a group of genes, each of them with *I_g_* isoforms, where *g* = 1,…,|*G*|. At least one of the isoforms of each gene *g* in *G* must belong to the same cluster *c*, where *c*∈ {1,…,*C*} and *C* is the total number of clusters. Let **z** be the expression vector obtained after concatenating the expression vectors **y_i_** of each isoform *i* of every gene *g* = 1,…,|*G*|. For the sake of simplicity, let us assume that |*G*|=2, *I_g_*=2 ∀ *<ι>∀∀g</i>*, and consequently *C*=2. In this case, vector **z** will contain 4*N* elements, where *N* is the total number of cells in the data (*N*=241 in our data) and will be the response variable in our regression model. We need to assess if **z** values follow the trend depicted in Figure 5B, that is, the average profile across cell types of the two isoforms in cluster 1 must be significantly different to the average profile of the two isoforms in cluster 2. In addition, the average profile of the two isoforms of gene 1 must not be different to the average profile of the two isoforms of gene 2. To identify groups of genes with these characteristics we proposed to fit the regression model in Equation 4 and select the group of tested genes as coDIU candidates when having a significant interaction between cluster and cell type effects, and a non-significant interaction between gene and cell type effects.

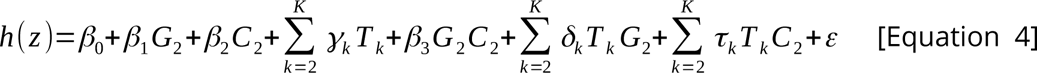

Where *G_2_* and *C_2_* are dummy variables indicating whether the expression value corresponds to gene or cluster 2 (value 1) or 1 (value 0), respectively, *T_k_* is a dummy variable which takes value 1 when the corresponding cell is assigned to cell type *k* (*k*=1,…,*K*) and 0 otherwis*e, βk, γk, δk* and *τ_k_* are the regression coefficients, *ε* represents the error term, and *h()* is the link function of the GLM (natural logarithm in this case).

We fitted the GLM model with the glm() function in the R-base package, and the *negative.binomial()* function in the *MASS* R package^94^, with θ = 10. To test the significance level of the *cluster*cell type* and *gene*cell type* interactions, we calculated type-II analysis-of-variance (ANOVA) tables for the model using a likelihood-ratio Χ^2^ test, implemented in the *Anova()* function from the *car* R package^95^, since we had an unbalanced design. *P*-values for each of the interactions were separately adjusted using the Benjamini & Hochberg correction. Gene pairs were considered positive for coDIU if FDR adjusted *p-*value < 0.05 for the *cluster*cell type* interaction and FDR adjusted *p-*value > 0.05 for the *gene*cell type* interaction. In other words, we required expression variance across cell types to be a function of the expression profile captured by the clustering, while imposing the additional limitation that aggregating expression by gene must make this effect undetectable. Given that all genes with clustered isoforms will form pairs with all potentially coDIU counterparts and be repeatedly tested, we considered genes to be positive for coDIU if they met the significance criteria in at least one of these pairwise tests.

### Functional analyses

The analyses in this manuscript are based on a long read-defined transcriptome which, after careful quality control and curation of the isoform models, was further annotated using IsoAnnotLite (https://isoannot.tappas.org/isoannot-lite/) to include positionally-defined functional features in the annotation (see Supplementary Note 1). *Functional features* are grouped in *functional categories* depending on the database from which the information was retrieved and on the biological functions performed by the features (comprehensive list in Supplementary Note 1). In this manner, we gathered sufficient information to couple our co-expression analyses with a biological readout. The specific analysis strategies used to this end are detailed below.

#### Functional Enrichment Analysis

In order to understand the functional properties of AS-regulated and co-regulated genes, we set out to characterize DIU and coDIU genes using different functional enrichment analysis approaches. In this manner, we intended to gain insight on functional features and categories showing significant overrepresentation in each of these two gene lists, in comparison to different backgrounds, i.e. lists of genes to compare to in order to detect enrichment.

In the case of DIU genes, we calculated enrichment relative to genes with multiple DE isoforms (total: 10,100) in order to discriminate the functional properties of genes regulated by alternative splicing, as opposed to those lacking differential usage of their isoforms. We considered all annotated functional categories and features, and applied tappAS Functional Enrichment Analysis (FEA), which relies on the *GOSeq* R package^96^. Briefly, the method performs an over-representation Fisher’s Exact test for each functional feature, considering the number of genes annotated with the feature in the tests and background lists. tappAS next corrects for multiple testing within each functional category by the Benjamini-Hochberg method, allowing multiple functional databases to be included or excluded from the analysis without influencing the number of significant features after p-value adjustment. Significant enrichment for the different tests was defined using a threshold of FDR<0.05.

For coDIU genes, we designed a different strategy in order to improve the statistical power of our functional enrichment analysis, aiming to compare functional properties between splicing regulation (DIU) and co-regulation (coDIU). As stated above, DIU regulation is best measured by using genes with DE isoforms as background. Intuitively, coDIU-regulated genes should then be characterized by comparing them to DIU genes. To accommodate these two test/background lists in a functional enrichment analysis without ignoring the overlap between the coDIU and DIU gene groups, we computed enrichment using a partially overlapping samples z-test via the *Prop.test()* function in the *Partiallyoverlapping* R package^61^. Specifically, we compared the proportion of coDIU genes containing each of the functional features (relative to DIU genes) with the proportion of DIU genes containing that same annotation (with respect to genes with DE isoforms). In other words, we tested whether the proportion of coDIU vs DIU genes including a given functional feature was significantly higher than that shown in the comparison between DIU and DE genes. We performed the analysis for features with more than 20 annotated genes, and subsequently corrected for multiple testing within functional categories using the Benjamini-Hochberg method. Functional features were considered to be present in a significantly higher proportion in coDIU genes when FDR<0.05.

#### Functional Diversity Analysis

To obtain insight into the functional changes generated as a consequence of DIU and coDIU, we again used the tappAS tool for the functional analysis of alternative splicing^60^. In particular, we first applied tappAS’ Functional Diversity Analysis (FDA) module (see main text Figure 6A). Briefly, FDA performs a within-gene comparison of all the isoforms included in the analysis, aiming to detect whether they present variation in the inclusion of a functional feature. In FDA, variation can be positional, i.e. one or more of the gene’s isoforms present a change in the genomic coordinates defining the feature, or be defined by presence/absence, i.e. at least one of the isoforms lacks a feature that is present in the rest. As a result, FDA provides analyzed genes with a label for each of the feature categories included in the transcriptome’s functional annotation file, flagging them as *varying* if at least one of the isoforms presents variation in a feature from that category, or *not varying* if no changes are detected. For more details on FDA, see the Methods section in de la Fuente *et al.*^60^.

We run both positional and presence/absence FDA for three gene sets: 1) genes with multiple DE isoforms (total: 3,172), 2) DIU genes (total: 2,577) and 3) coDIU genes (total: 2,049). Next, for each of these gene sets, we computed the proportion of *varying* genes detected for each functional category. Varying proportions were calculated relative to the total number of genes including annotations from the category, instead of considering all genes in the set. In this manner, we avoided underestimating variation rates for categories that were less represented in the functional annotation file. In order to check whether any of these gene sets presented a significantly higher mean proportion of varying genes across categories, we performed a paired *t*-test for each combination of gene set pairs: DIU vs multiple DE, coDIU vs multiple DE, and coDIU vs DIU. In this analysis, we considered functional categories to be the individuals under evaluation, while the proportion of varying genes calculated for each category in the two tested sets constituted the paired observations. As a result, we obtained three p-values per FDA analysis type, i.e. presence/absence and positional variation.

To better understand the functional readout that can be obtained using the *acorde* pipeline, we analyzed a subset of the coDIU gene network, namely three clusters showing related isoform co-expression patterns: neuron-specific expression (cluster 8), oligodendrocyte-specific expression (cluster 13) and expression in both neural and oligodendrocyte cell types (cluster 19). In total, 160 coDIU genes showed co-expressed isoforms between at least two of these clusters. To characterize functional variation among the clusters, we used positional/presence FDA (see above) and ID-level FDA. ID-level FDA is also included in tappAS^60^ and provides a within-feature summary of FDA results. In other words, ID-level FDA ultimately reports the number of *varying* and *not varying* genes detected for each feature ID included in a given functional category. In this case, *varying* status obeys a similar criterion to the one described above, i.e. genes in which at least one isoform shows differential inclusion/exclusion of the feature. Since each functional category may include several features, ID-level FDA provides a complementary view to that of FDA, allowing users to inspect which particular features are more frequently changing as a result of the category-level functional variation reported in FDA. For more details on ID-level FDA, see the Methods section in de la Fuente *et al*.^60^.

## Author contributions

AAL developed and evaluated computational methods, implemented and documented R package, obtained long read transcriptome, analyzed single-cell data, interpreted results and wrote the manuscript. PS implemented IsoAnnotLite, refined and performed functional annotation strategy. ST conducted statistical method development, refined the manuscript and supervised the study. AC conceived the study, supervised all analysis approaches and methods developed, and refined the manuscript.

## Acknowledgements

This work has been funded by NIH grant R21HG011280 and by the Spanish Ministry of Science grants BIO2015–71658 and BES-2016-076994.

## Competing interests

The authors declare no competing interests.

---

**Supplementary Figure 1:**
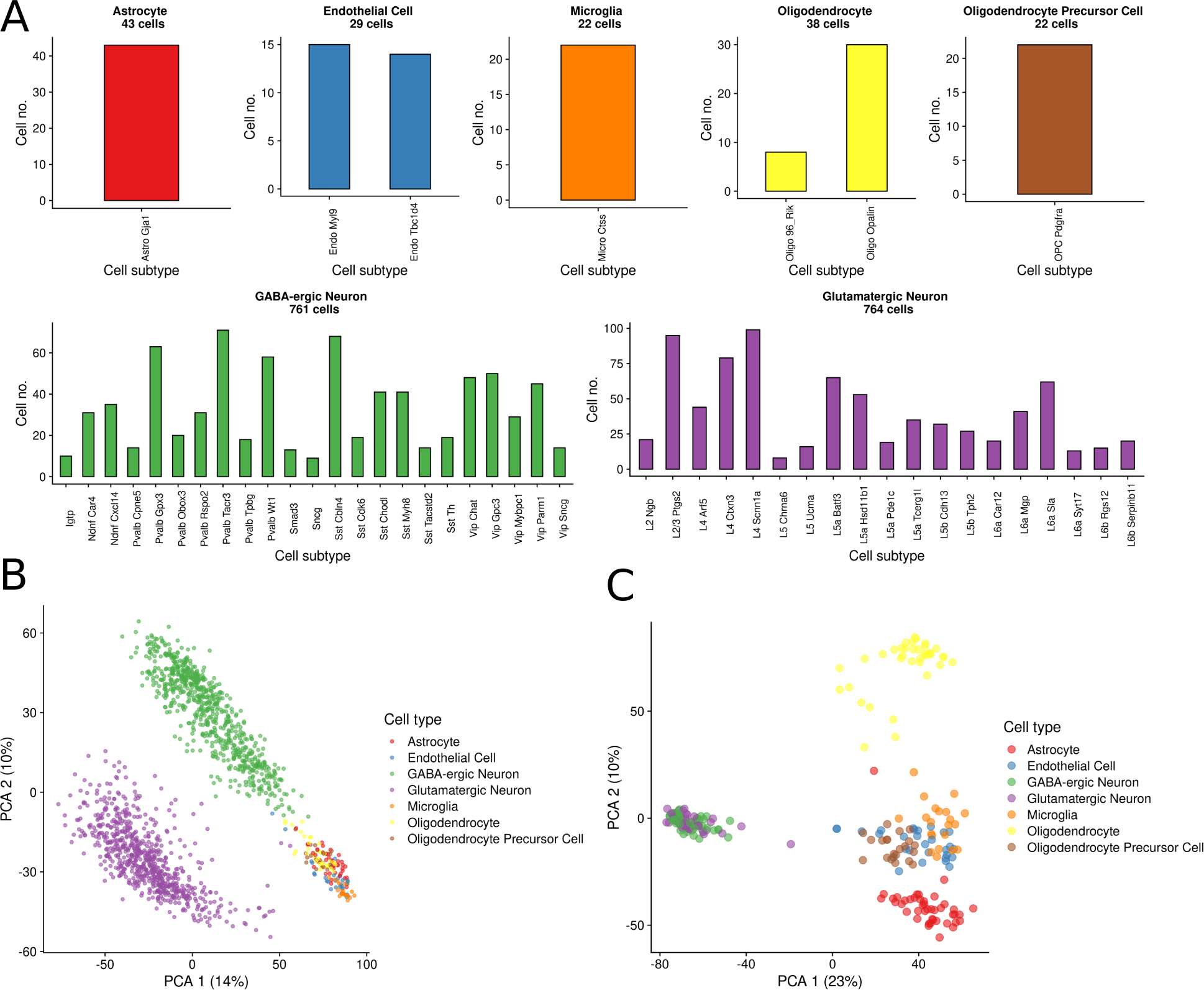
Tasic et al. mouse neural dataset overview. A) Number of cells assigned to each cell type and subtype by Tasic et al. (post-QC, 1591 cells). B) Principal Component Analysis of the full dataset, components 1 and 2. C) Principal Component Analysis of data after balancing cell number by randomly sampling 50 GABA and 50 glutamatergic neural cells (242 total cells).

**Supplementary Figure 2:**
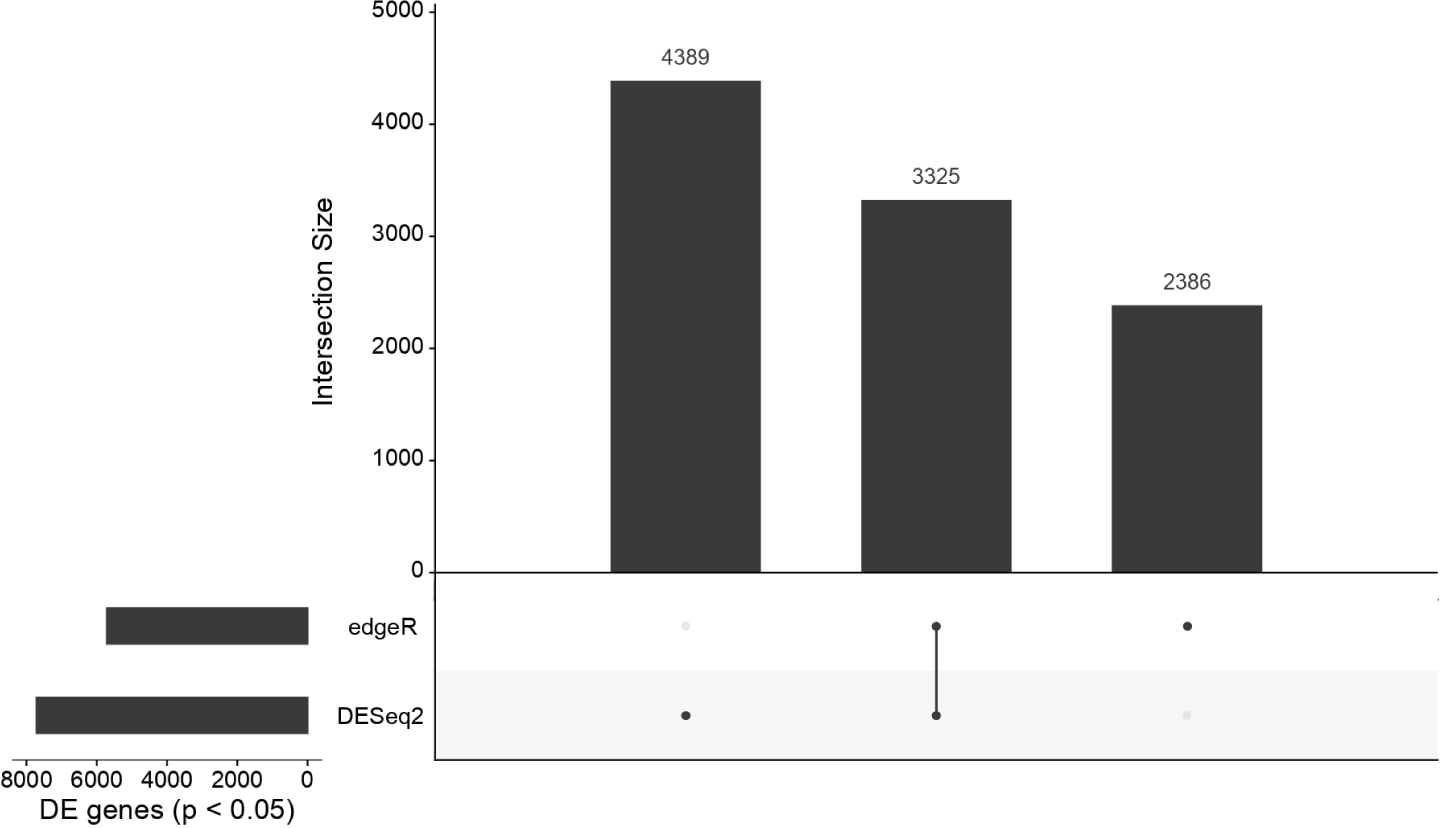
upset plot of isoform-level Differential Expression (DE) analysis results. DE isoforms common and uniquely-detected by edgeR and DESeq2 are represented by vertical bar height. Analyzed sets (unique isoforms or intersection) are indicated by the dots below. Total DE isoforms detected by each method (adjusted *p-*value < 0.05) is represented by horizontal bar height.

**Supplementary Figure 3.**
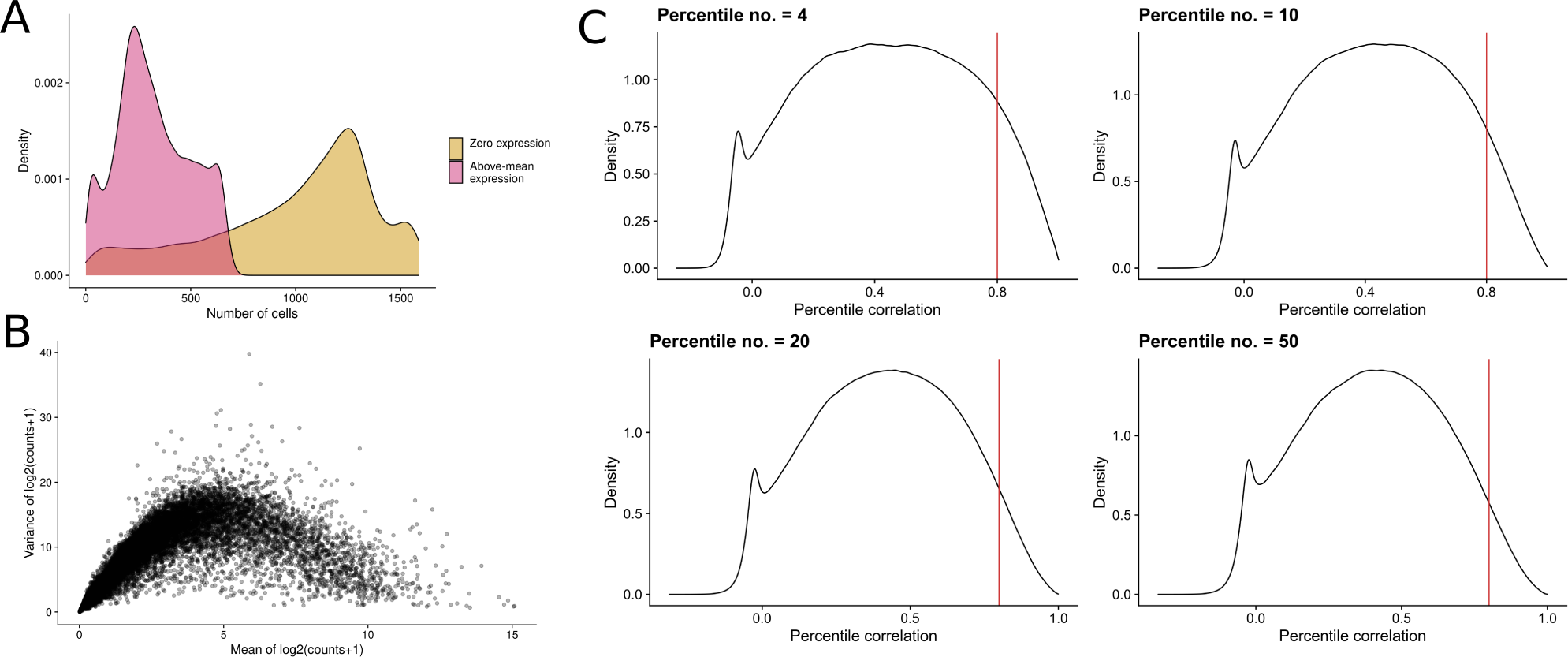
A) Heterogeneity and sparsity in the Tasic et al. dataset. x-axis shows number of cells per isoform showing zero expression (yellow) and expression higher than the isoform mean (pink), y-axis shows density. B) Dataset mean-variance relationship. Dots represent transcript isoforms, x-axis represents the isoform mean and y-axis correspond to isoform expression variance (log2counts). C) Percentile correlation density distributions obtained after using different number of percentiles to summarize cell type-level expression.

**Supplementary Figure 4:**
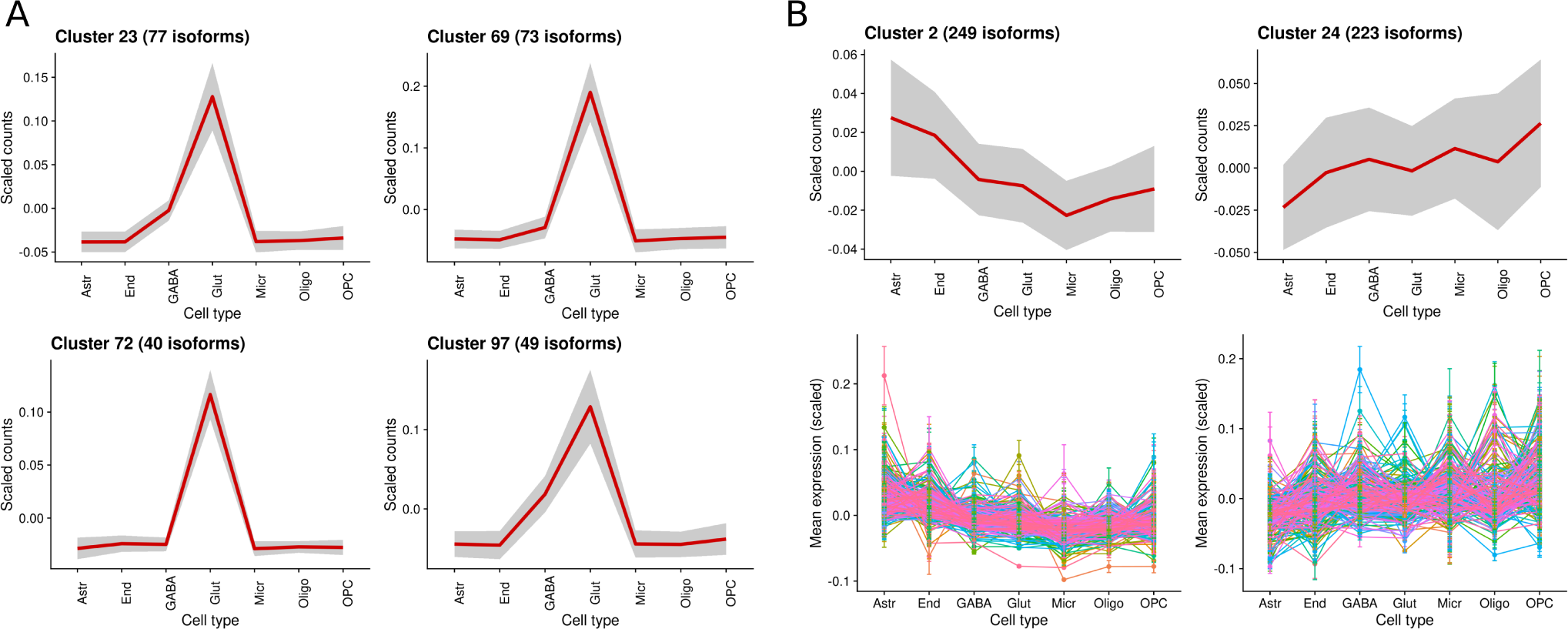
cluster refinement examples. A) Example of four redundant clusters, i.e. clusters representing the same expression profiles, merged by profile similarity. B) Example of noisy clusters, i.e. clusters grouping isoforms with highly dissimilar expression patterns across cell types, whose isoforms were re-assigned to other clusters.

**Supplementary Figure 5:**
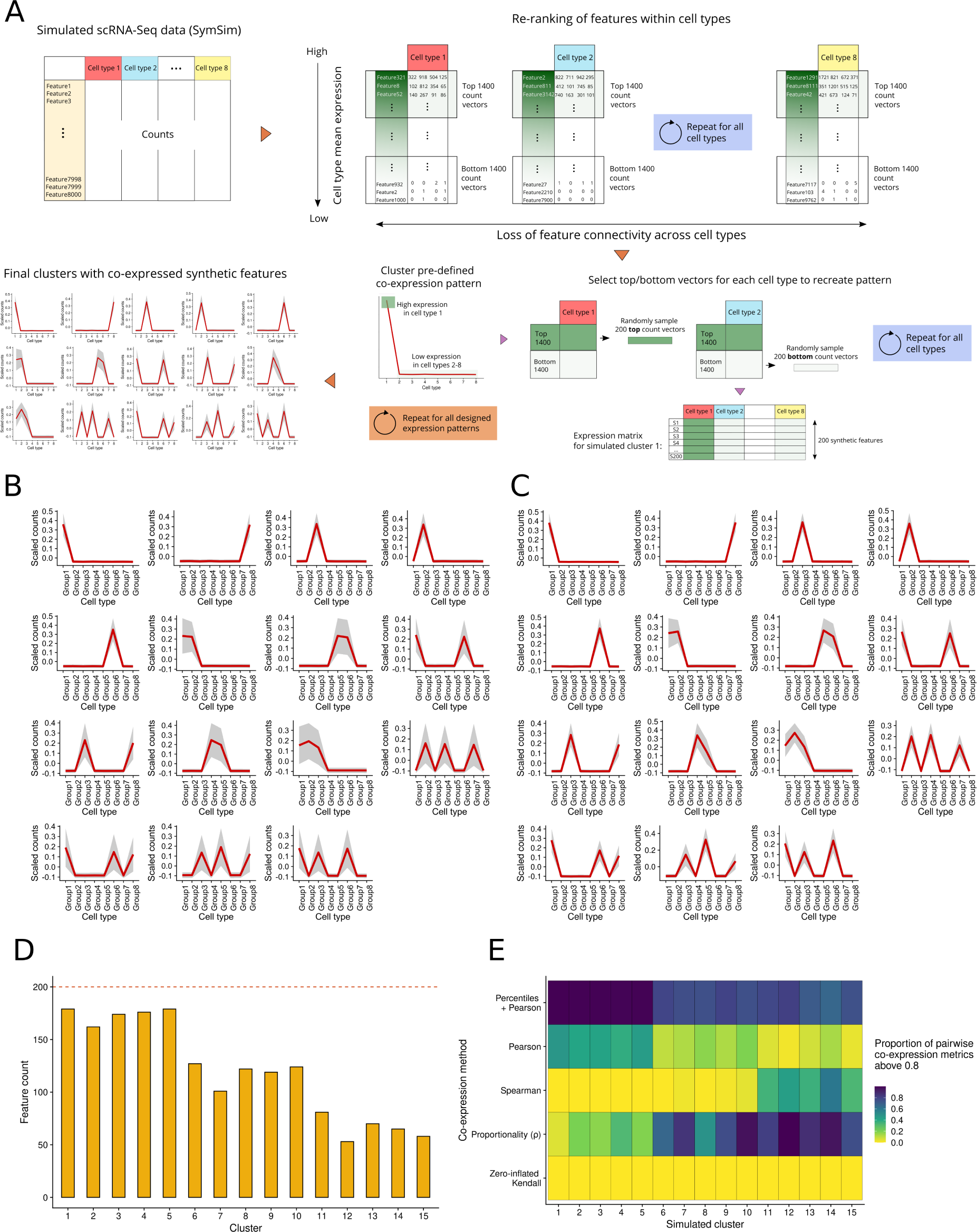
simulated data characterization. A) Single-cell co-expression pattern simulation strategy. After simulating single-cell RNA-seq data with SymSim, features are re-ranked within each cell type, using the simulated expression values to generate groups of co-expressed transcripts. Simulated clusters were generated according to 15 pre-defined patterns. B) and C) Expression profile of the 15 simulated clusters obtained, before (B) and after (C) filtering synthetic features showing dissimilarity with the simulated profile. Cell-level mean expression (scaled, see Methods) is computed for all simulated transcripts in the cluster and then aggregated as the global cell type mean, represented by the red line. Grey area corresponds to cell type mean ± standard deviation. D) Number of transcripts in simulated clusters after filtering to ensure cluster profile consistency. E) Heatmap representing the proportion of high co-expression values (correlation or proportionality > 0.8), computed for all pairs of synthetic transcripts within each simulated cluster. Darker colors indicate successful recapitulation of simulated co-expression relationships by the evaluated co-expression metrics.

**Supplementary Figure 6:**
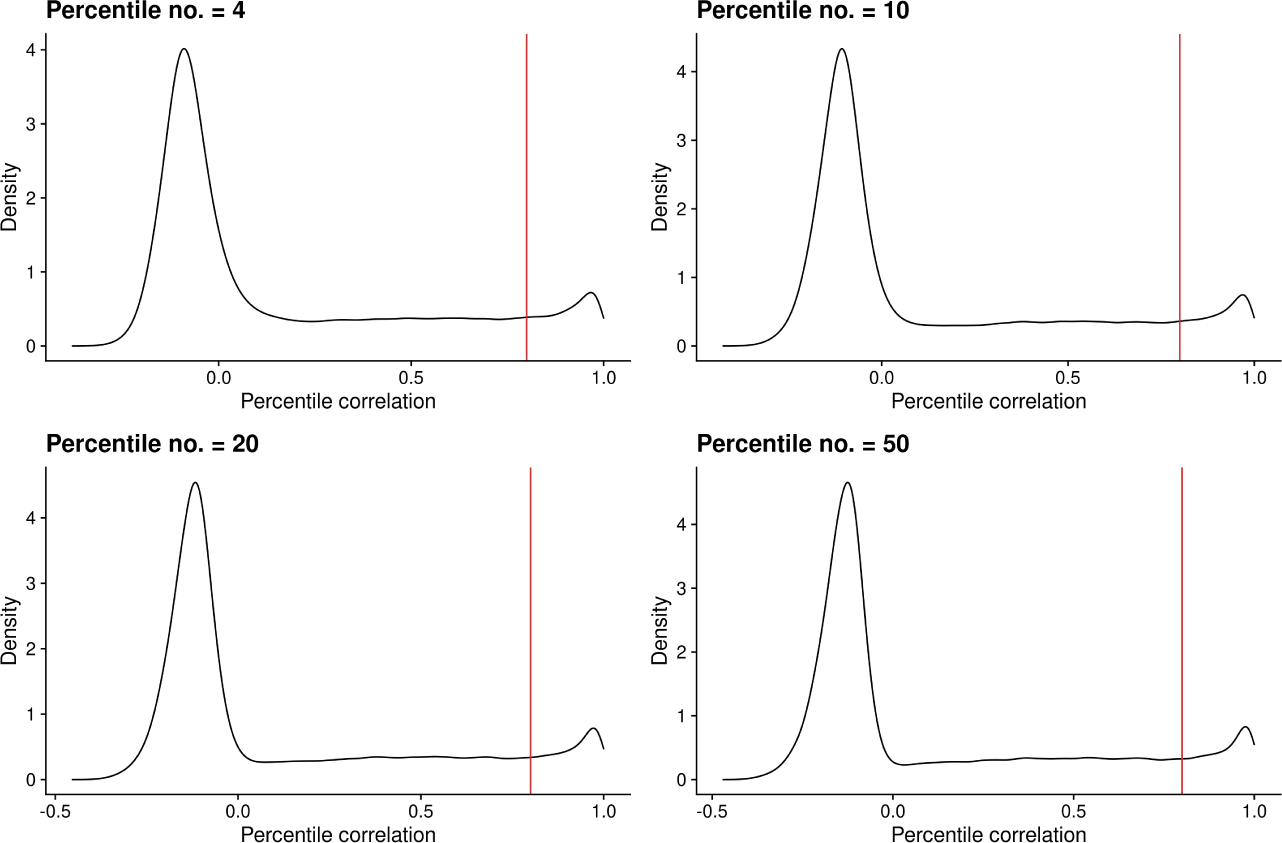
effect of percentile number in simulated data. Percentile correlation density distributions obtained after using different number of percentiles to summarize cell type-level expression.

**Supplementary Figure 7:**
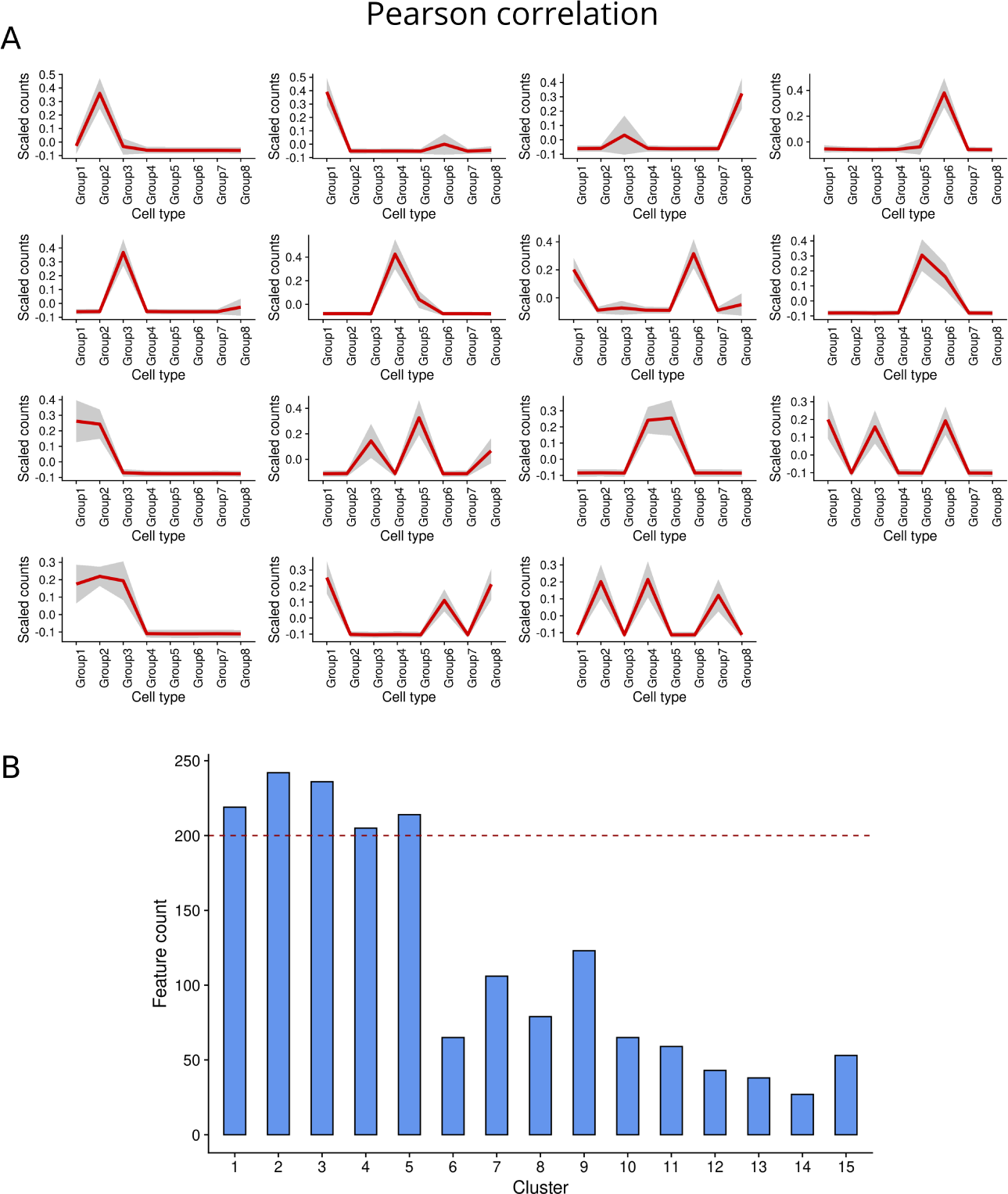
clustering of simulated single-cell data using Pearson correlation as distance. A) Mean profiles of generated clusters. Cell-level mean expression (scaled, see Methods) was computed for all simulated transcripts in the cluster and then aggregated as the global cell type mean, represented by the red line. Grey area corresponds to cell type mean ± standard deviation. B) Number of simulated transcripts per cluster.

**Supplementary Figure 8:**
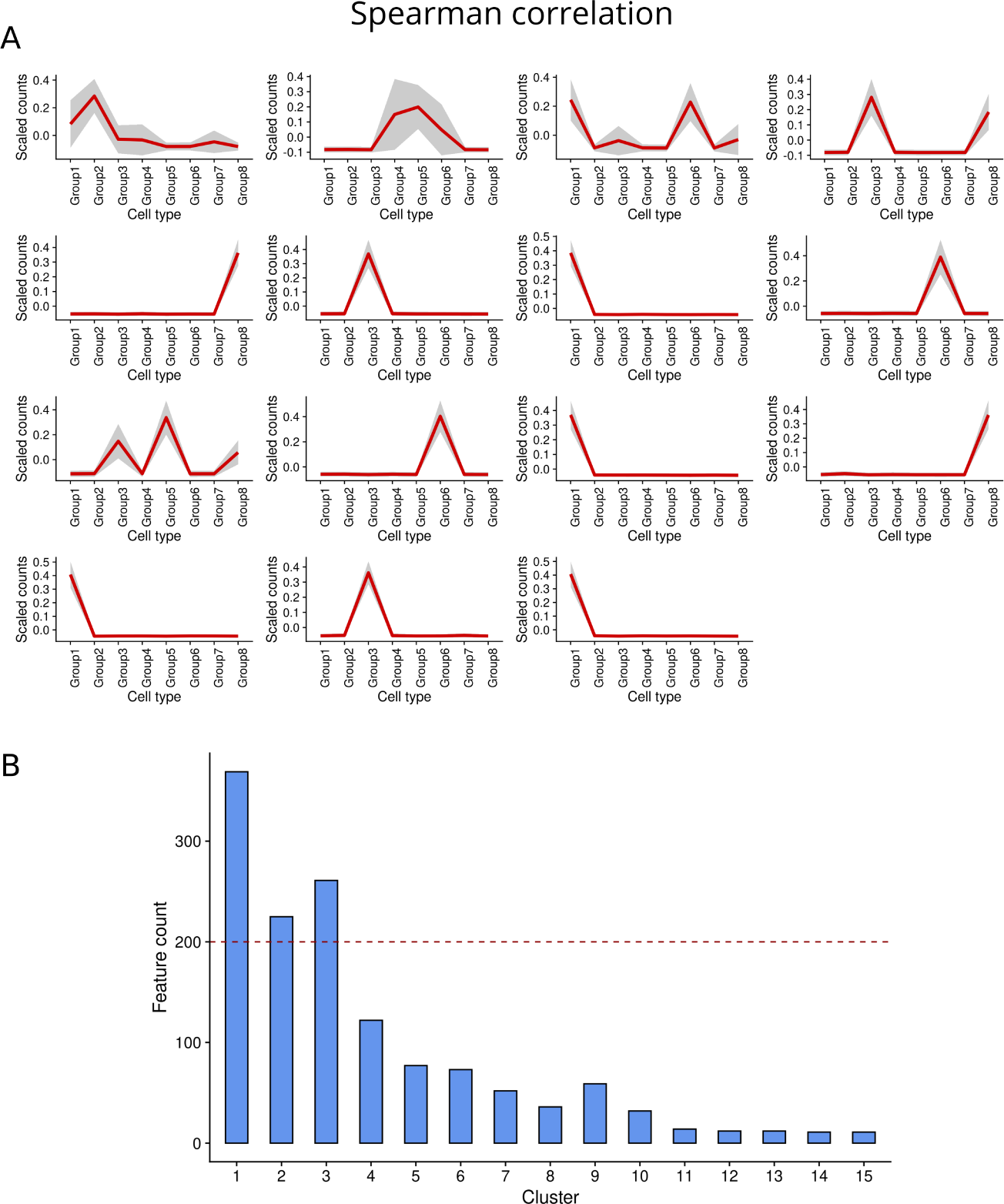
clustering of simulated single-cell data using Spearman correlation as distance. A) Mean profiles of generated clusters. Cell-level mean expression (scaled, see Methods) was computed for all simulated transcripts in the cluster and then aggregated as the global cell type mean, represented by the red line. Grey area corresponds to cell type mean ± standard deviation. B) Number of simulated transcripts per cluster.

**Supplementary Figure 9:**
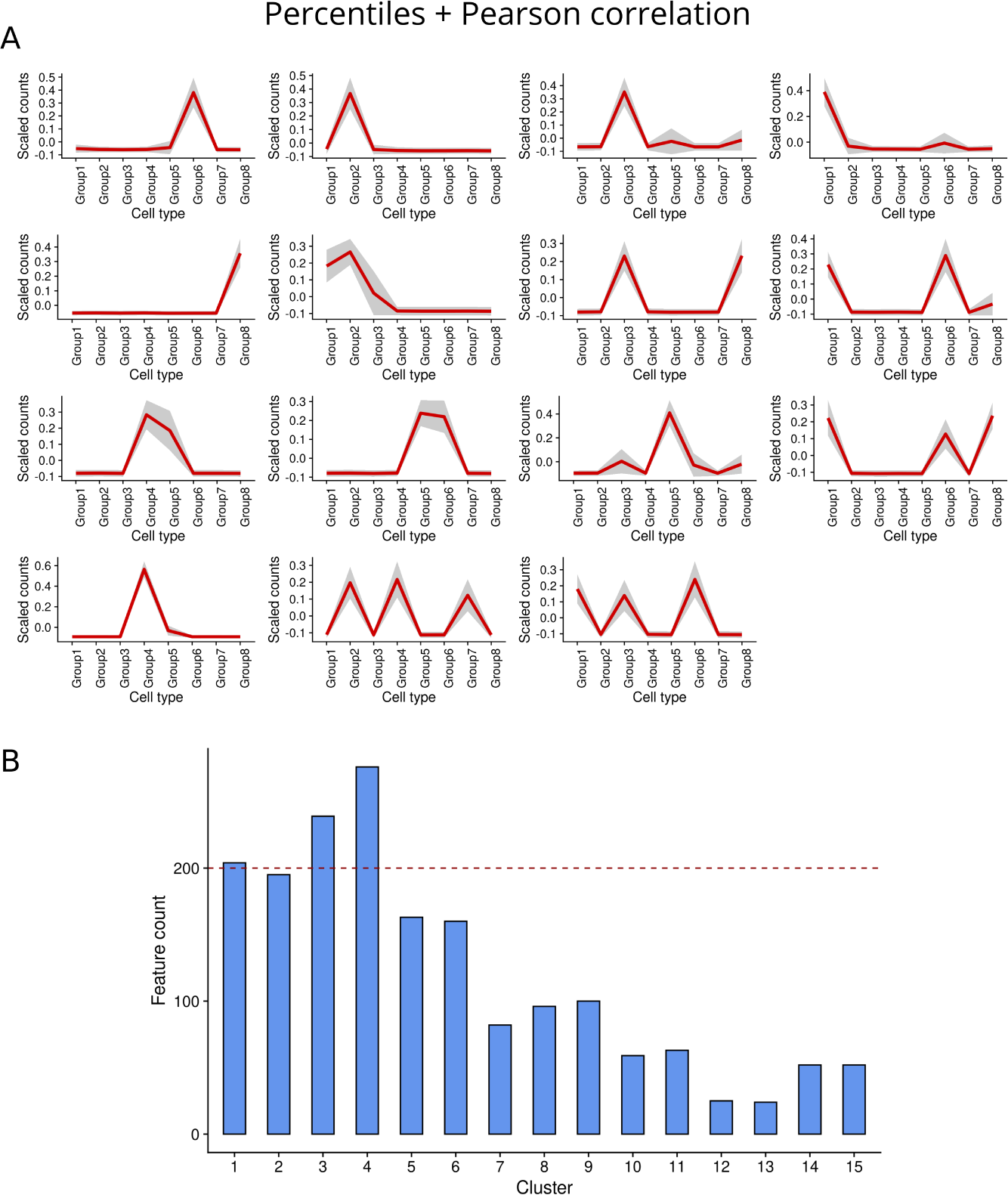
clustering of simulated single-cell data using percentiles + Pearson correlation as distance. A) Mean profiles of generated clusters. Cell-level mean expression (scaled, see Methods) was computed for all simulated transcripts in the cluster and then aggregated as the global cell type mean, represented by the red line. Grey area corresponds to cell type mean ± standard deviation. B) Number of simulated transcripts per cluster.

**Supplementary Figure 10:**
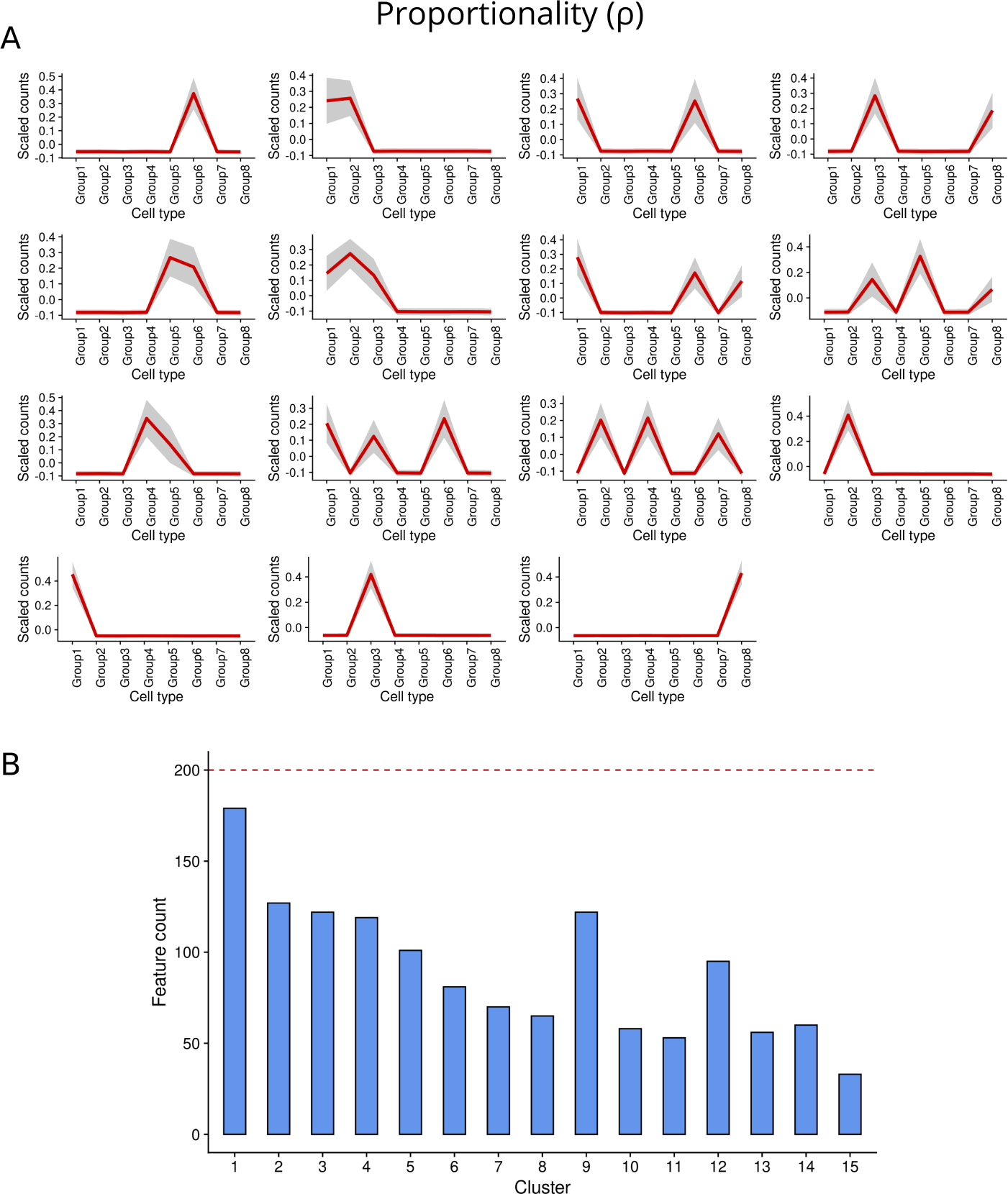
clustering of simulated single-cell data using proportionality (ρ) as distance. A) Mean profiles of generated clusters. Cell-level mean expression (scaled, see Methods) was computed for all simulated transcripts in the cluster and then aggregated as the global cell type mean, represented by the red line. Grey area corresponds to cell type mean ± standard deviation. B) Number of simulated transcripts per cluster.

**Supplementary Figure 11:**
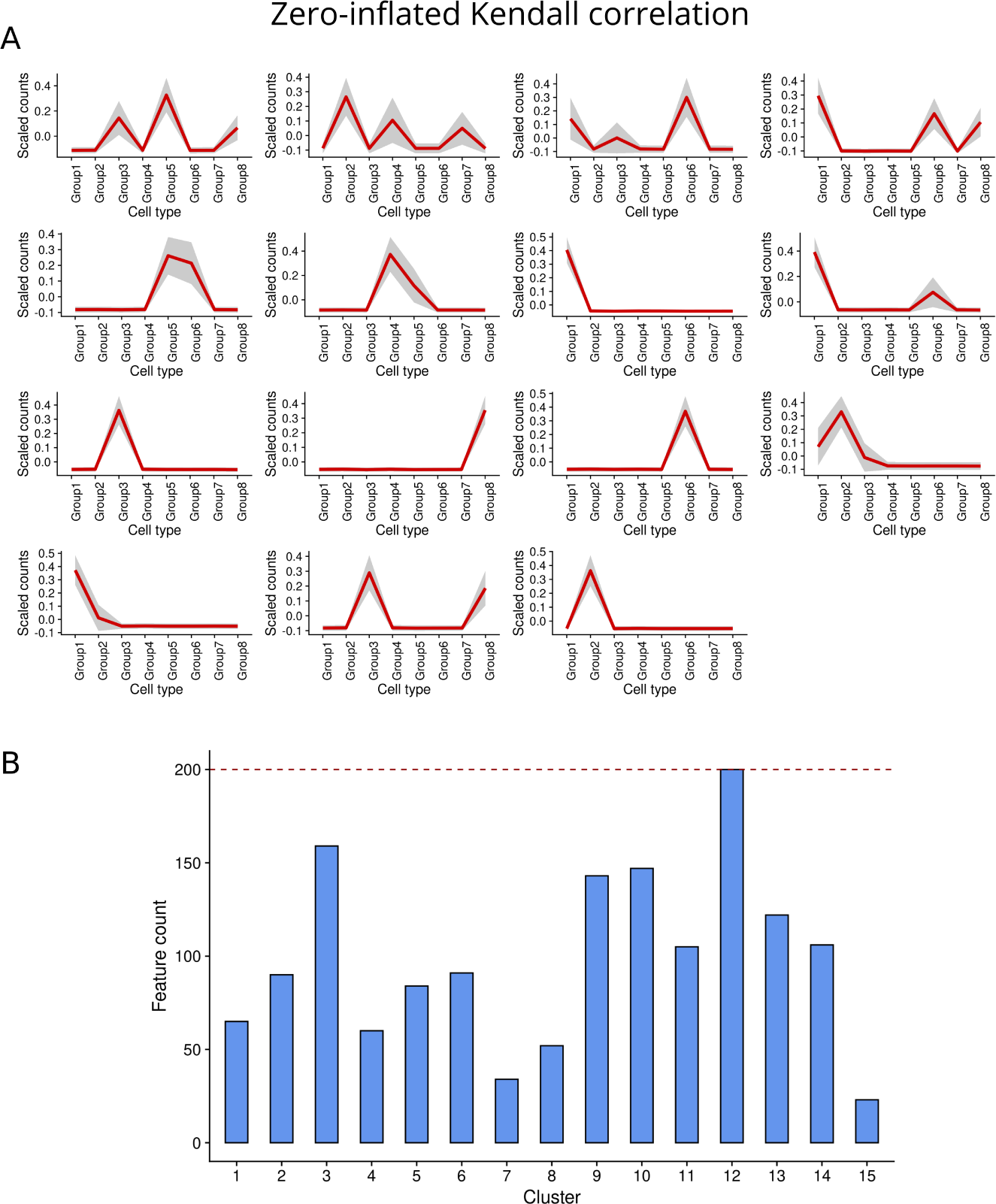
clustering of simulated single-cell data using zero-inflated Kendall correlation as distance. A) Mean profiles of generated clusters. Cell-level mean expression (scaled, see Methods) was computed for all simulated transcripts in the cluster and then aggregated as the global cell type mean, represented by the red line. Grey area corresponds to cell type mean ± standard deviation. B) Number of simulated transcripts per cluster

**Supplementary Figure 12:**
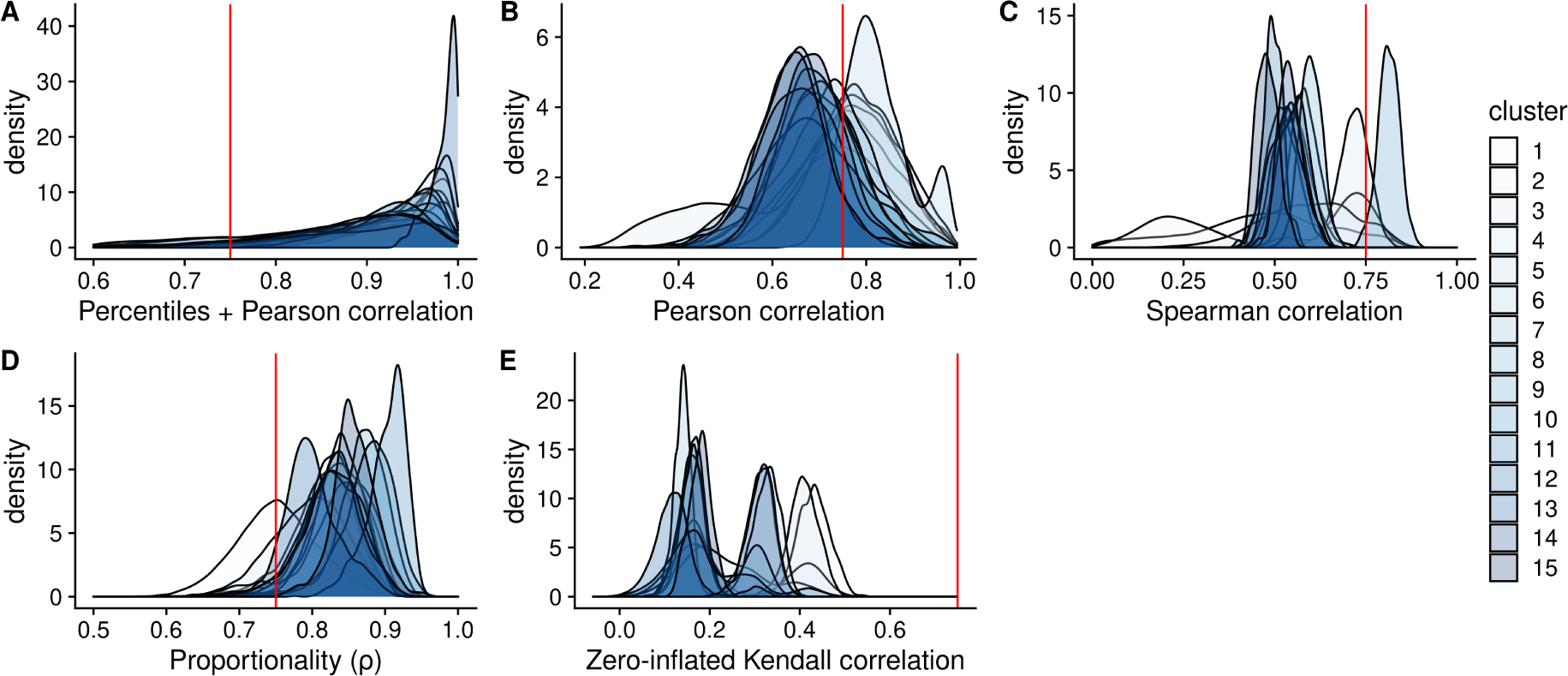
density distributions of co-expression values between isoforms in clusters obtained from simulated data. Red line indicates the 0.75 threshold value. Results shown for the clusters generated using each evaluated metric: A) Percentiles + Pearson correlation. B) Pearson correlation. C) Spearman correlation. D) Proportionality (ρ). E) Zero-inflated Kendall correlation.

**Supplementary Figure 13:**
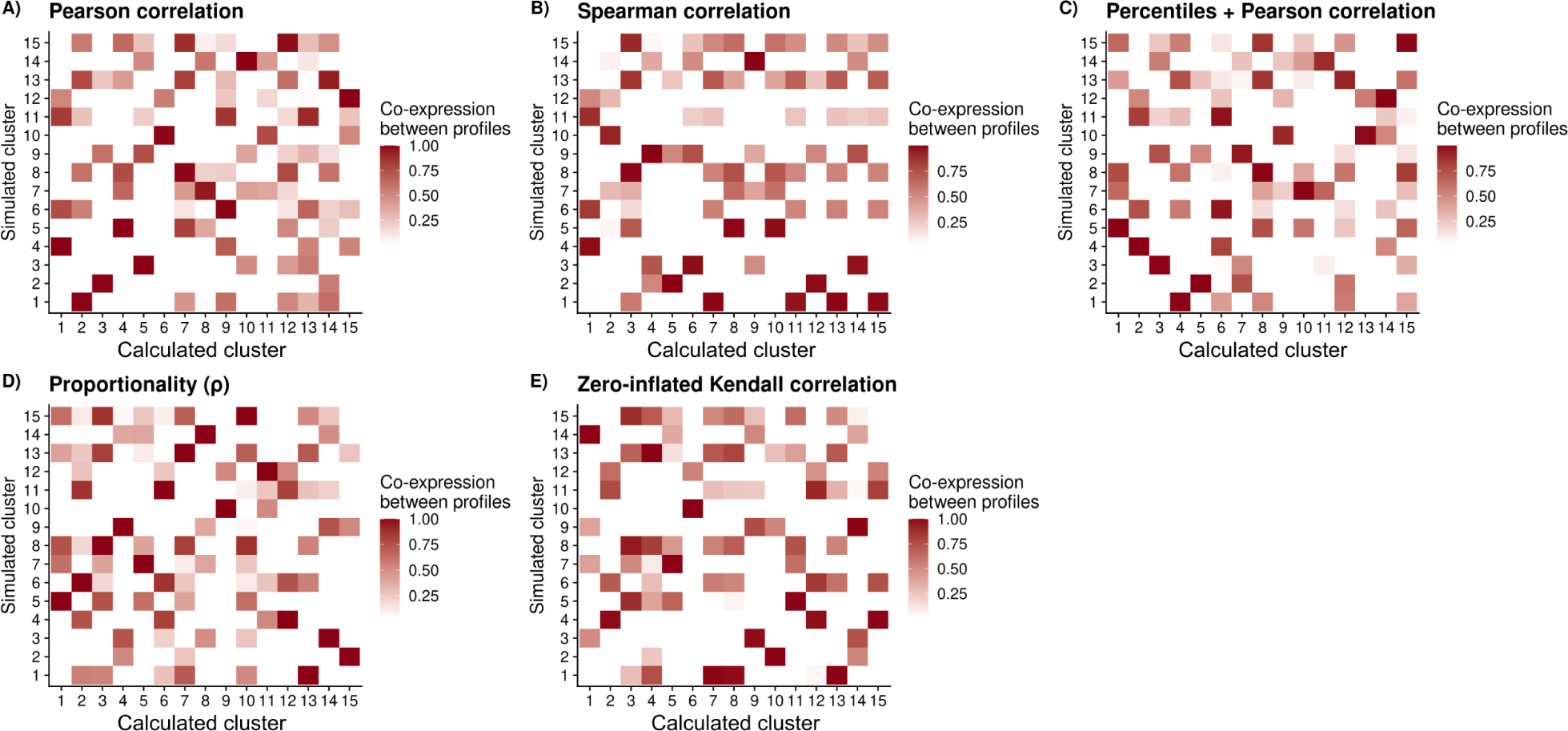
Pearson correlation between mean expression profiles of calculated and simulated clusters. Results shown for the clusters generated using each evaluated metric: A) Pearson correlation. B) Spearman correlation. C) Percentiles + Pearson correlation. D) Proportionality (ρ). E) Zero-inflated Kendall correlation. Calculated clusters show no unique match among simulated clusters and were therefore paired with the simulated cluster showing the highest mean expression profile correlation.

**Supplementary Figure 14:**
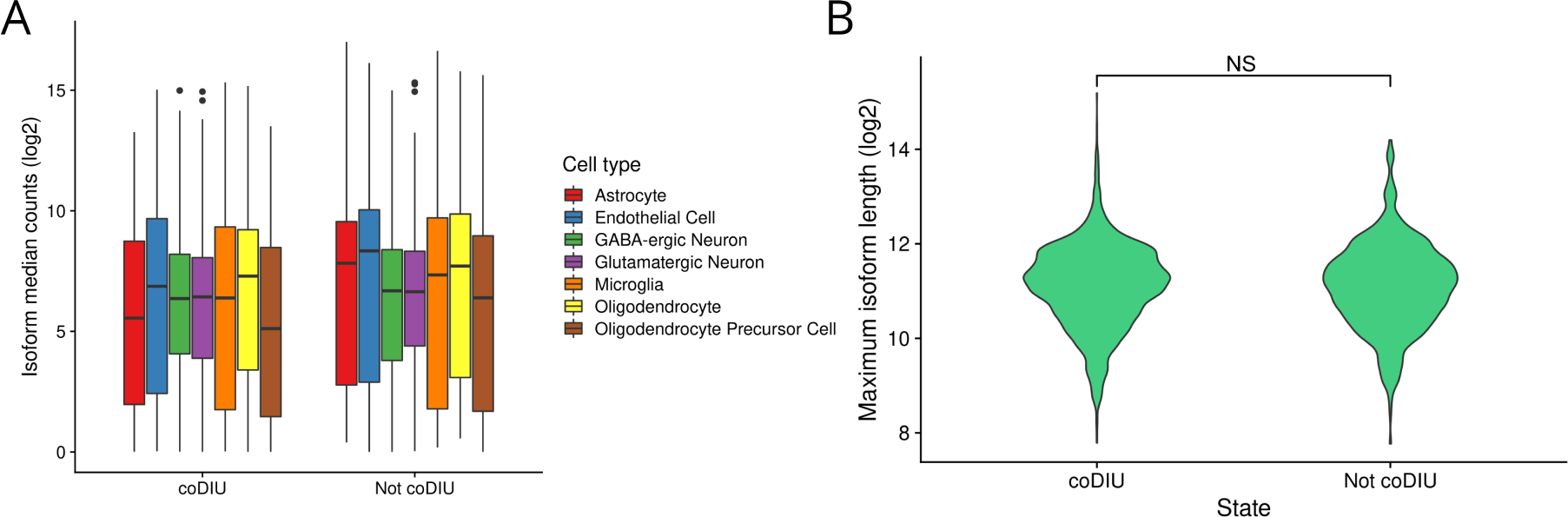
evaluation of expression and length biases on coDIU genes. A) Boxplot of cell type-level median counts (log2) for isoforms belonging to coDIU and not coDIU (i.e. solely DIU) genes. Dots represent expression outliers. B) Violin plot of isoform length (log2) of longest isoform in coDIU and not coDIU (i.e. solely DIU) genes. No significant differences were found between them (NS: not significant, Wilcoxon test).

**Supplementary Figure 15:**
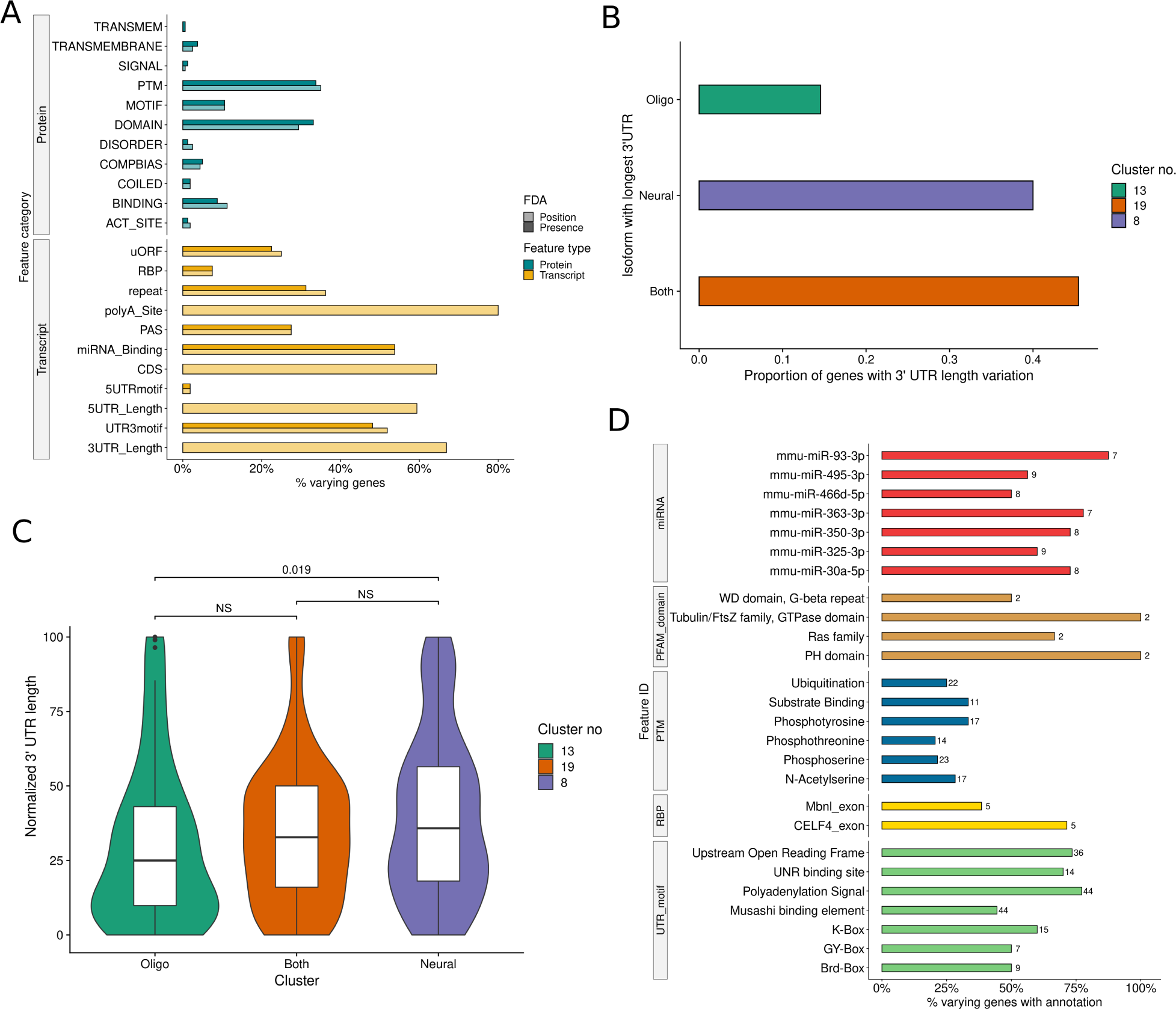
functional analysis of the 160 coDIU genes detected across neural
-oligodendrocyte clusters (no. 8, 13 and 19). A) Functional Diversity Analysis (FDA) results. y-x: functional annotation categories. x-axis: percentage of genes including at least one *varying* feature from a given functional category. B) Proportion of coDIU genes with 3’UTR variation across isoforms that have their longest 3’UTR isoform in each of the three analyzed clusters. C) Violin + boxplot of normalized 3’UTR lengths for each of the three neural-oligodendrocyte clusters. Normalized lengths are computed by dividing each individual isoform’s 3’UTR length by the sum of 3’UTR lengths of all the gene’s isoforms. Significance levels for the comparison of the three groups are indicated above the corresponding braces (*p-*value, Wilcoxon test). D) ID-level FDA results for features in highly-varying functional categories. Percentage of genes containing the feature that show functional variation across isoforms (x-axis) is shown for the most frequently annotated features in each category. Total coDIU genes with feature indicated by the bar label.

**Supplementary Figure 16:**
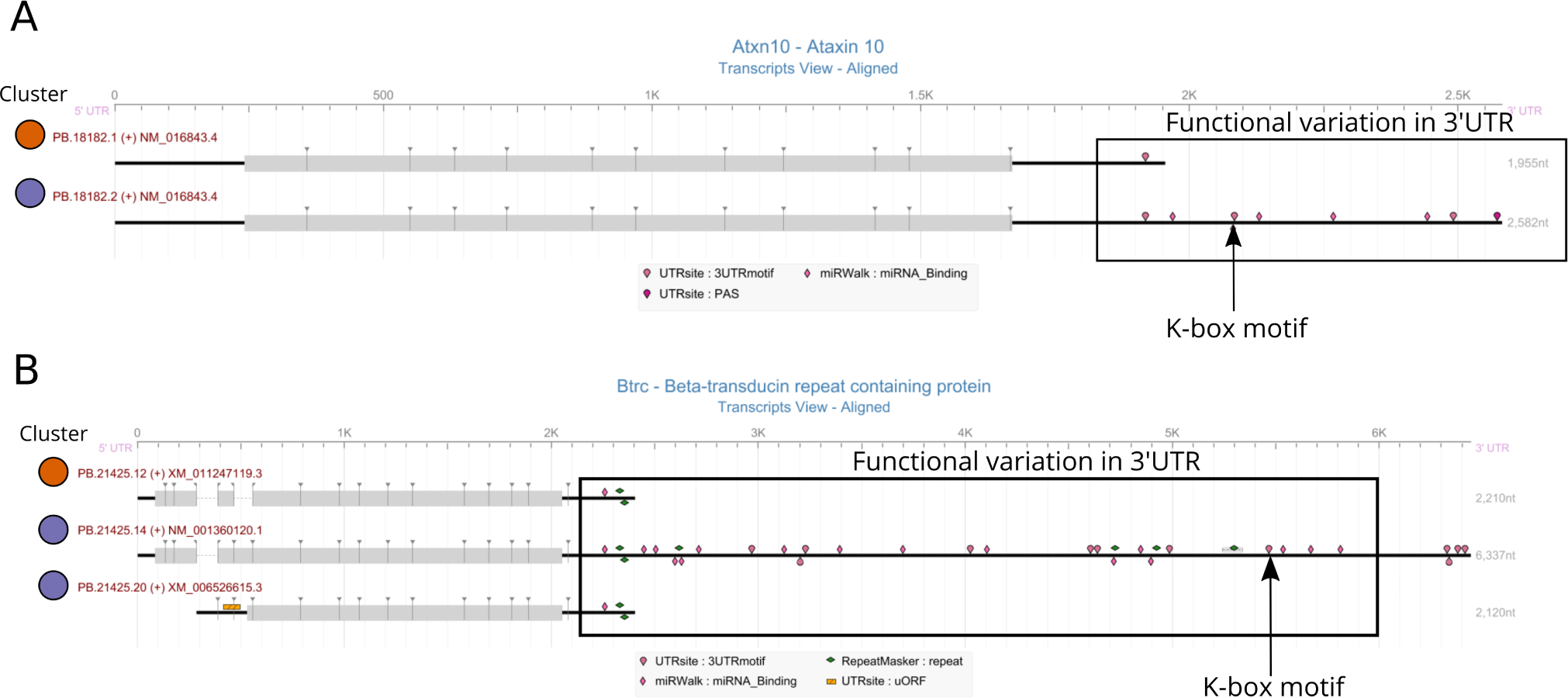
tappAS view of transcript functional annotation for A) *Atxn10* and B) *Btrc* isoforms. Cluster assignments for each isoform are indicated by dot color.

## Notes

### Competing Interest Statement

The authors have declared no competing interest.

