## Supplementary notes for "*Acorde*: unraveling functionally-interpretable networks of isoform co-usage from single cell data"

### Supplementary Note:

### Obtaining a high-quality, long read-defined transcriptome

As a part of our study, we have combined bulk PacBio data for neural transcriptome definition and single-cell short-read RNA-Seq data for cell-level isoform quantification. The purpose of this Supplementary Note is to detail the strategy that was used build a high-quality, long read-defined transcriptome using PacBio data, including analytical decisions made during pre-processing, quality control with SQANTI3 and curation using both SQANTI's filtering tools and additional information provided by external supporting data (Figure 1).

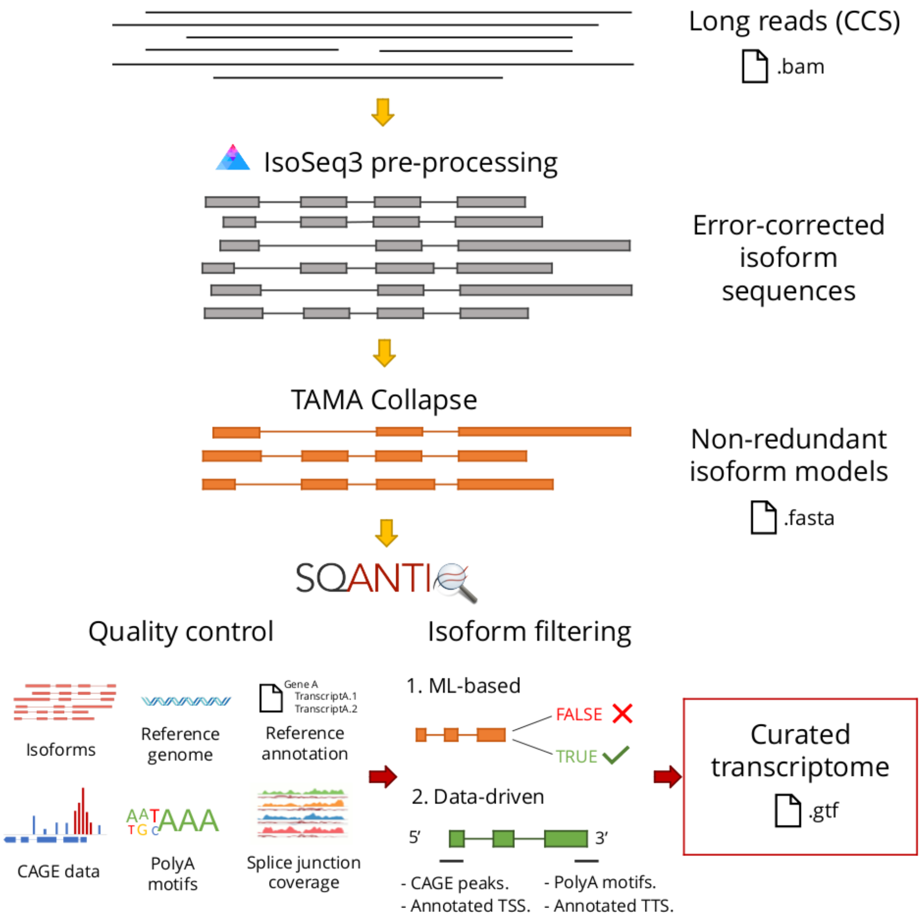


**Figure 1:** pipeline overview of long read data pre-processing and quality control using SQANTI3.

#### *Long read data and pre-processing*

Long-read datasets from mouse hippocampus and cortex[1] were downloaded from ENCODE accessions ENCSR214HSG and ENCSR340GWV, respectively. To obtain a single mouse neural transcriptome, reads from both samples and their replicates (2 replicates/sample) were pooled and pre-processed using the IsoSeq3 pipeline (https://github.com/PacificBiosciences/IsoSeq) (including inter-read correction step, i.e. polishing), with default parameters. After pre-processing, we obtained 178507 isoforms from 20307 genes and characterized the transcriptome using SQANTI3 QC (https://github.com/ConesaLab/SQANTI3) isoform categories (Figure 2A).

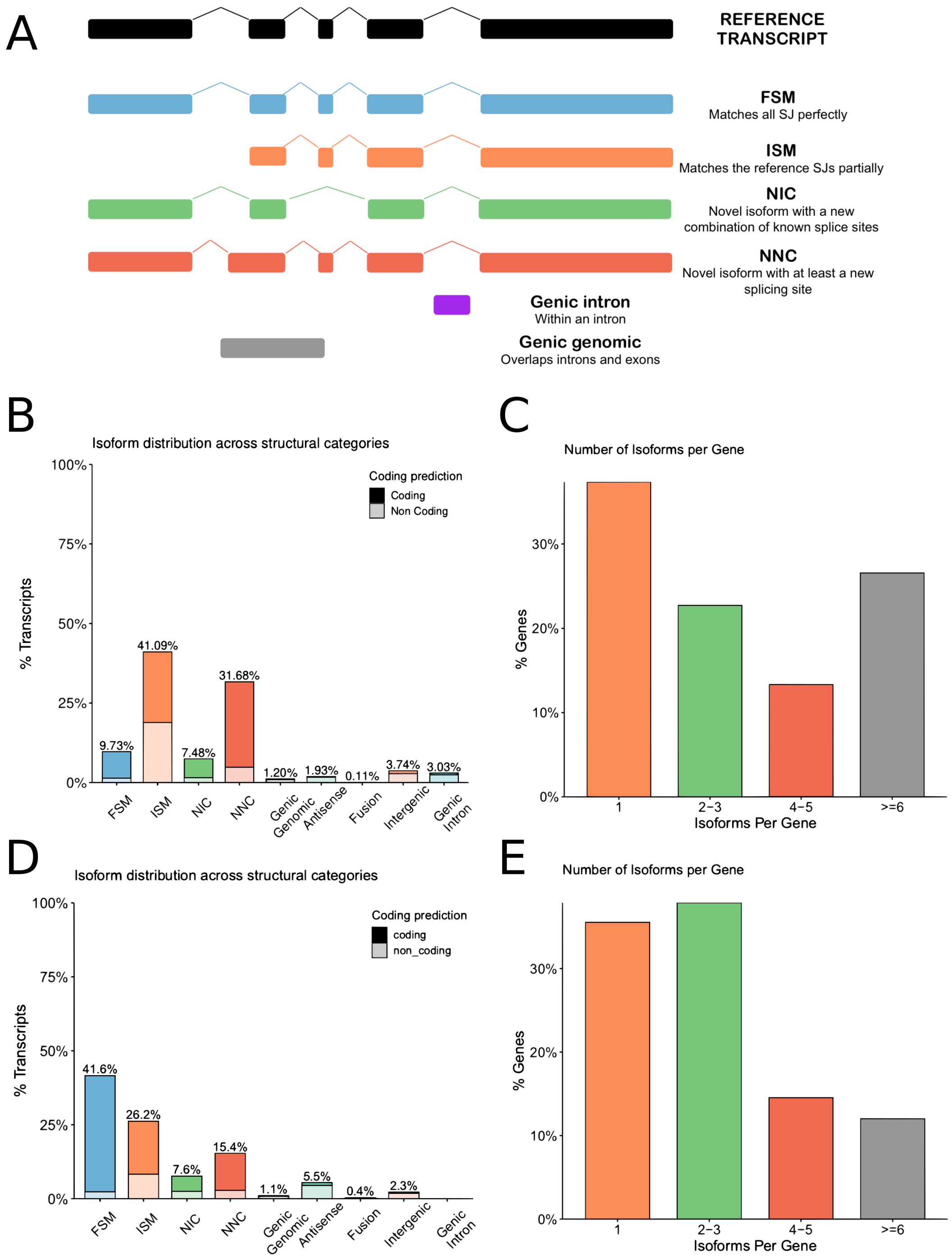

**Figure 2: transcriptome overview after pre-processing vs after collapse.** A) Description of the main SQANTI3 structural categories (obtained from https://github.com/ConesaLab/SQANTI3). B) Isoform distribution across structural categories for the mouse neural transcriptome after IsoSeq3 pre-processing. C) Number of isoforms per gene after IsoSeq3 pre-processing. C) Isoform distribution across categories after collapse with TAMA. D) Number of isoforms per gene after collapse with TAMA.

Given the dataset's enrichment in Incomplete Splice Matches (ISM, Figure 2B), and to avoid an overestimation of the number of isoforms per gene (Figure 2C), we collapsed redundant isoform models using TAMACollapse[2] (https://github.com/GenomeRIK/tama/wiki/Tama-Collapse). In this manner, we intended to mitigate transcript-level redundancies by merging isoform models that differ due to library preparation and sequencing artifacts, but are very likely to have been generated from the same transcript in the sample. We run TAMACollapse with the following parameters: -c 95 -i 85 -x no_cap -a 10 -m 10 -z 10. This reduced the transcriptome to 93698 isoforms. Collapsing removed a big proportion of ISM isoforms (Figure 2D) and successfully reduced the complexity of our transcriptome (Figure 2E).

*Supporting data for long read transcriptome quality control and curation*

To perform the quality control of our transcriptome, we run SQANTI3 (https://github.com/ConesaLab/SQANTI3) on the collapsed isoform sequences, using the mouse reference genome (GRCm38.p6) and the RefSeq[3] transcriptome (global release 96, annotation release 108, accessed December 2019). To perform a comprehensive evaluation and subsequent curation of the quality of our isoforms, we also provided supporting data to SQANTI3, namely CAGE peak data, polyA motif sequences and short read splice junction coverage data (Figure 1).

Mouse CAGE peak data were obtained from the FANTOM5 database[4], and used to curate Transcription Start Sites (TSS) for our isoforms. Consistent with previous reports stating that human and mouse polyA motif sequences show high levels of conservation[5,6], we supplied a ranked list of common polyA motif sequences from human (provided with SQANTI3) to curate Transcription Termination Sites (TTS). Finally, to obtain short-read coverage information, and given the lack of matching Illumina data for these long read datasets, we sampled 20 cells from each of the 7 broad cell types in the Tasic *et al.* dataset[7] and pooled all Illumina reads to generate a pseudo-bulk. We mapped this cell pool to the long read-generated transcriptome (genome version GRCm38.p6) using STAR[8] and the following parameters:

*--outFilterType BySJout --outFilterMultimapNmax 20 --alignSJoverhangMin 8 --outFilterMismatchNmax 999 --outFilterMismatchNoverReadLmax 0.04 --alignIntronMin 20 --alignIntronMax 1000.*

*Transcriptome curation and characterization*

In order to curate the long read transcriptome, we defined a strategy for isoform filtering to ensure the quality and realiability of the transcript models (Figure 1). First, we applied SQANTI's random forest classifier[9] (henceforth refered to as ML filter) to perform quality filtering of Novel In Catalog (NIC) and Novel Not in Catalog (NNC) isoforms as well as minority categories such as Antisense, Genic Genomic and Fusion. Next, the above-detailed sources of supplementary data were integrated with information from RefSeq's reference transcriptome (RefSeq release 96, September 2019, genome: GRCm38.p6, annotation: GCF_000001635.26) to remove artifacts belonging to the Full Splice Match (FSM) and Incomplete Splice Match (ISM) categories. To better ilustrate the rational behind this two-step strategy, Figure 3 provides an overview of how each filter modifies transcriptome properties based on SQANTI3 structural categories.

During the first stage of curation, SQANTI's ML filter used 11 isoform-level features, including junction coverage information, to detect false positive isoforms. Of note, this tool was originally designed to detect transcript models that are likely originated by RT-switching, intra-primming and the accumulation of sequencing errors around junctions. Isoforms flagged as artifacts by the classifier were directly removed, reducing our transcriptome from 93698 to 74148 isoforms. In this process, ~40% NIC and ~80% of NNC isoforms were classified as artifacts (Figure 4A), whereas isoforms from other minority categories were subject to equally stringent filtering (Figure 4B). NIC and NNC isoforms labeled as artifacts by SQANTI's ML filter were enriched in low coverage junctions (Figure 4C) and novel splice sites, with the unfiltered transcriptome showing accumulation of non-canonical sites in the NNC category (Figure 4D). While low support of was especially frequent among novel junctions, isoforms with low-coverage novel splice sites were successfully removed by SQANTI's ML filter (Figure 4E-F).

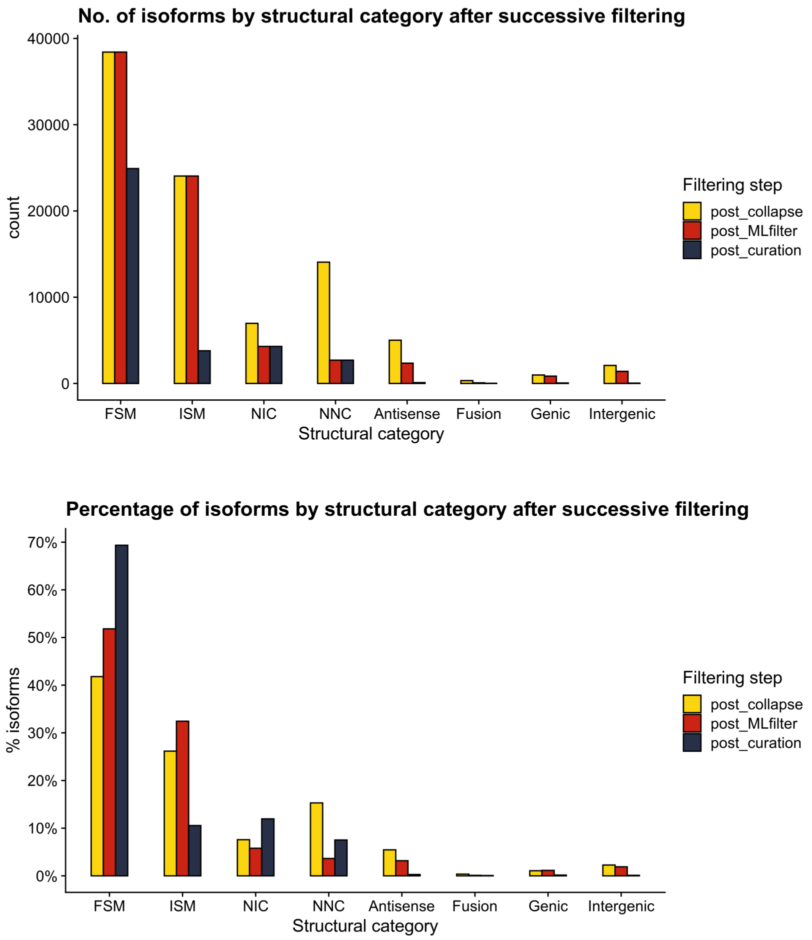


**Figure 3:** Number and proportion of isoforms within each SQANTI3 structural category in the post-collapse transcriptome and after performing each filtering step.

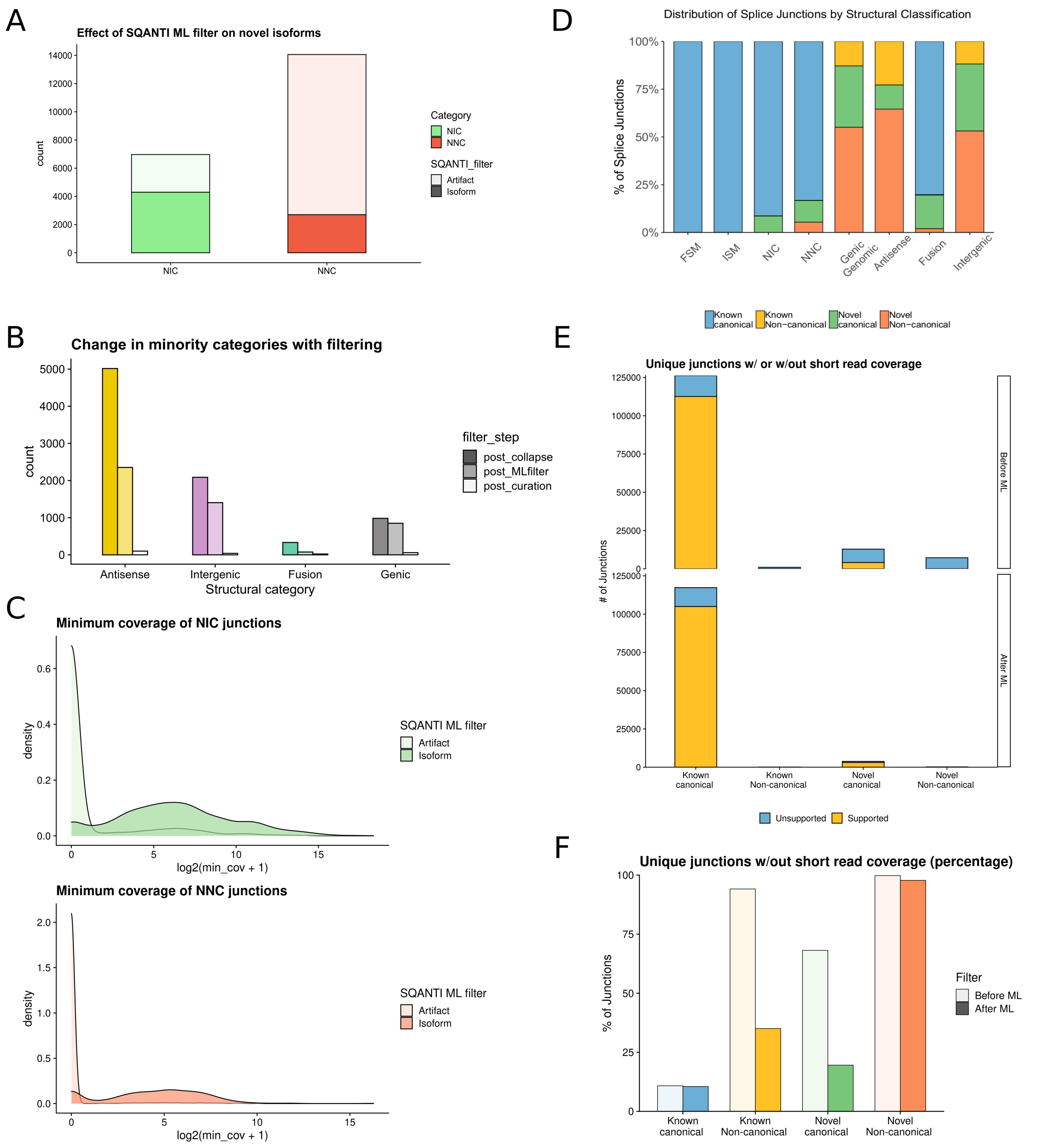

**Figure 4: characterization of SQANTI's ML filter.** A) Number of NIC and NNC isoformsm flagged as artifacts or isoforms by SQANTI's ML filter. B) Number of isoforms removed after each filtering step in minority structural categories. C) Density distribution of the minimum short-read coverage of junctions, shown for NIC (left) and NNC (rigth) isoforms and stratified by SQANTI's ML filter result. D) Splice junction type (%) present in each SQANTI structural category. E) Number of unique junctions with or without short read coverage for each type of splice junction, before (upper side) and after (lower side) applying SQANTI's ML filter. F) Percentage of unique junctions with or without short read coverage for each type of splice junction, before and after the ML filter.

Next, we established filtering criteria for ISM and FSM isoforms. Transcripts in these two categories are used as true positives in SQANTI's ML filter, and therefore will be preserved without evaluating their reliability. However, while these categories solely present know (and therefore validated) junctions, they often incorporate 3' and 5' end variability, which can lead to the detection of multiple FSM and/or ISM isoforms per reference transcript, i.e. isoforms for whom all junctions have been previously reported by the reference annotation, but differing in their start and/or end positions. We named this phenomenon "redundancy". Importantly, redundancy can be the result of true TSS and TTS diversity or stem from library preparation artifacts such as RNA degradation or intra-priming, resulting in false positive novel TSS and TTS. In our transcriptome, high levels of redundancy were observed for both FSM (Figure 5A) and ISM (Figure 5B). To mitigate this, the amount of redundancy that can be confidently preserved was evaluated using CAGE peak data (5' end) and polyA motif information (3' end).

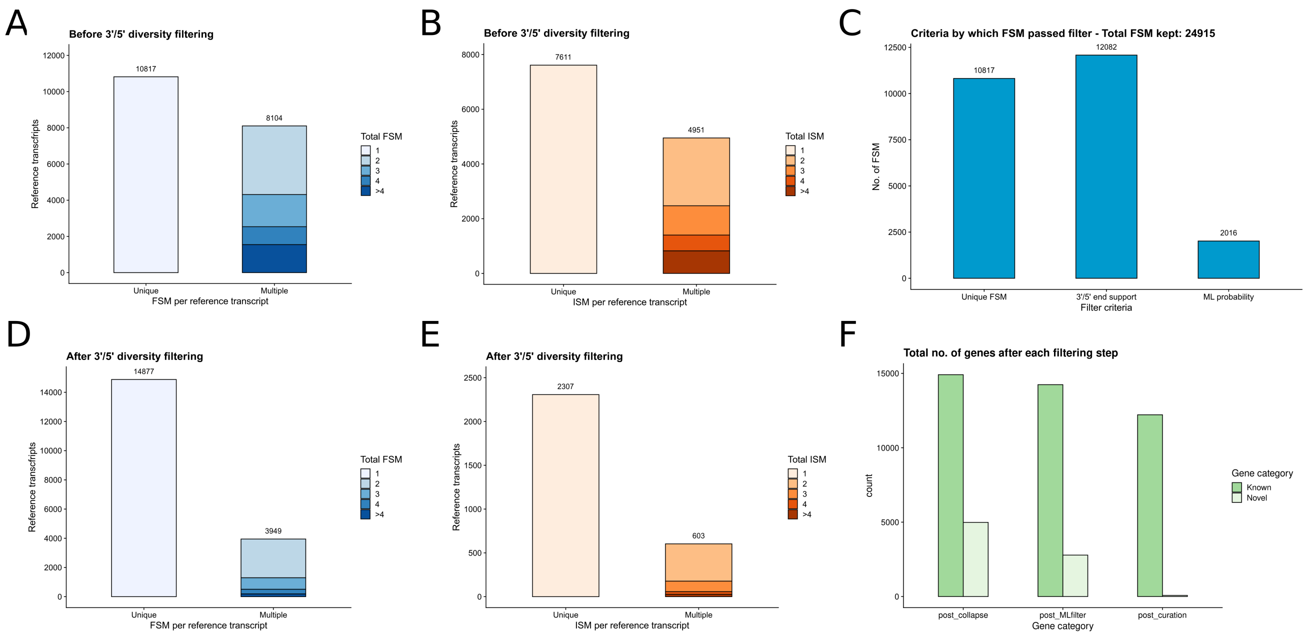


**Figure 5: effect of multiple-criteria filtering on the long read transcriptome.** A) and B) Redundancy levels for FSM (A) and ISM (B) before filtering based on CAGE and polyA data. C) FSM count stratified by the criteria used to included the isoform in the transcriptome. D) and E) Redundancy levels for FSM (D) and ISM (E) after filtering. F) Number of known and novel genes after collapse, and after each filtering step.

For FSM isoforms, we evaluated the suitability of preserving an isoform in the transcriptome according to three criteria, in the following order of priority: a) detection of a reference-compatible FSM transcript, in cases where there is no redundancy and even if no support is found; b) transcripts with support for TSS and TTS to preserve only reliable FSM, if there is redundancy; c) selection of the FSM that best matches the reference transcript, when there was redundancy but no 3'/5' end support for any of the FSM found. For ISM, given that missing junctions from matching reference transcripts increases the probability of false epositive TSS/TTS, we only applied criteria (b). These rules and their effect on the transcriptome will be described in more detail below.

First, transcripts that constituted the only matching FSM of a given reference transcript, i.e. unique-FSM cases, were automatically preserved, independently of CAGE/polyA support of their TSS and TTS. In this manner, we prioritize evidence of detection of junction combinations present in reference transcripts over TSS/TTS definition accuracy. Next, we evaluated TSS and TTS end positions to discriminate false positive isoforms in multiple-FSM cases, i.e. those where multiple FSM per reference transcript were detected. Regarding the 5' end, we kept isoforms whose TSS was included within or situated up to 50bp downstream of a CAGE peak, to account for degradation, and whose 3' end region included a polyA motif no further than 50bp upstream of the TTS. Alternatively, isoforms lacking CAGE or polyA support were included in the transcriptome if their TSS matched or was situated up to 50bp downstream of an annotated TSS, and their TTS also matched or was within ±50bp of an annotated TTS. In this manner, we consider both CAGE/polyA support and the reference transcriptome for curation, and include all possible combinations of these two sources in our transcriptome. For instance, an isoform will be included if it contains a novel TSS supported by CAGE, and its TTS is close enough to that of the reference transcript. ISM isoforms were similarly filtered according to polyA/CAGE and annotated TSS or TTS evidence. Finally, for multiple-FSM cases where no FSM met the TSS/TTS requirements, we devised a strategy to select the best match among detected isoforms. In this manner, and similarly to the unique FSM scenario, we prioritize evidence of detection of a reference-annotated combination of junctions. We preserved the FSM transcript with the highest random forest probability score, as output by the ML filter in SQANTI.

As a result of curation, 35700 isoforms from 12183 genes were preserved. All in all, this filtering strategy not only managed to enrich the transcriptome in FSM (24915 total FSM, ~67%, Figure 3), but also pointed out the importance of combining multiple criteria to prevent losing relevant FSM transcripts that would have been lost based solely on 3'/5' end support, with ~50% of FSM in our transcriptome being preserved based on unique FSM or best match criteria (Figure 5C). Furthermore, filtering effectively reduced redundancy in both FSM (Figure 5D) and ISM (Figure 5E), simultaneously retaining some multiple matches for both categories. Part of the 3'/5' diversity in the long read-defined transcriptome could therefore be attributed to true TSS and TTS, however, no isoforms from novel genes were preserved, given their lack of 3'/5' end support (Figure 5F) and thus our inability to tell true isoforms apart from artifacts.

Since we combined different data sources for filtering, we decided to inspect whether supporting data or agreement with the reference annotation (or both) had been the reasons for inclusion in the transcriptome. In doing so, we found only moderate levels of agreement between CAGE and polyA data and the reference annotation, highlighting the importance of supporting data for transcriptome curation. In the case of FSM, more than 50% of validated TSS and TTS could only be confirmed using CAGE and polyA motif data (Figure 6A), and this percentage was even higher for some ISM sub-categories (Figure 6B). For instance, internal fragments (missing junctions at both the 3' and 5' ends in comparison to the matching reference transcript) could only be validated using CAGE or polyA information. All in all, these analyses suggest that reference transcripts may contain multiple, unannotated TSS and TTS that are can only be observed using long reads.

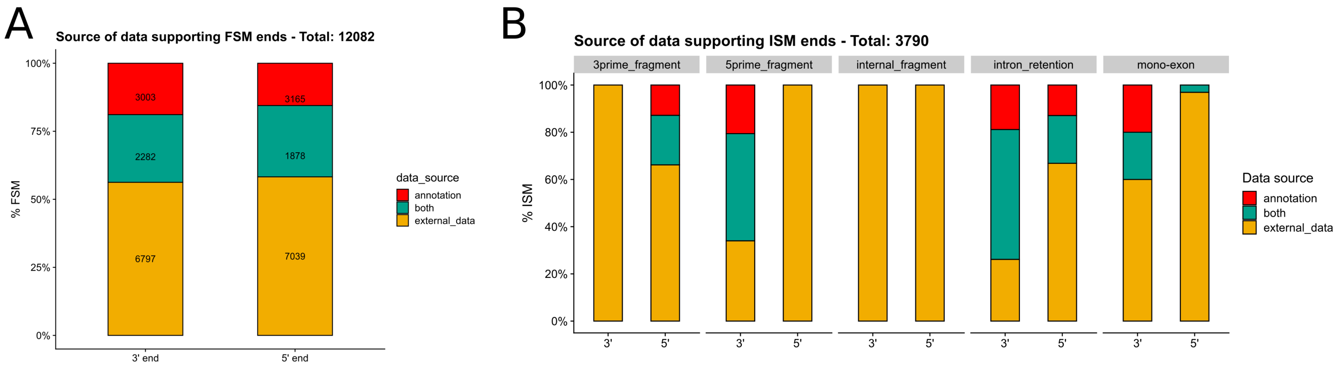


**Figure 6: source of data guiding isoform inclusion during end-support filtering.** A) Included FSM % supported by external (i.e. supporting) data, the reference annotation or both, for 3' and 5' end support. B) Included FSM % supported by each supporting data type, stratified by ISM sub-category.

In addition to the former, we performed a recovery process for genes for which multiple isoforms had been detected using long reads, but for which non of these long read-defined isoforms passed the TSS/TTS filtering. In these cases, reads compatible with multiple reference transcripts may have been found, even if long read data was not reliable enough for isoform model reconstruction. We focused on removed genes that had multiple FSM isoforms associated to two or more reference transcripts, and incorporated their associated reference transcripts from RefSeq to the transcriptome, while discarding the unsupported long read-defined isoforms. Consequently, we added 1286 missing isoforms and 509 genes for which there was evidence of alternative isoform expression, even if their isoforms could not be confidently defined using long reads. All in all, 36986 isoforms from 12692 genes were included in our final, curated transcriptome.

It should be noted that, for minority categories, checking for CAGE/polyA and annotation support resulted in the removal of practically all isoforms (Figure 4B). Given the difficulty in establishing the reliability of the remaining isoforms, or the relevance of CAGE/polyA data for this task, the few transcripts from categories that remained were excluded from the transcriptome.

#### *Functional annotation of long read-defined isoforms*

Recent studies by several research groups, including our own, have made a strong case for the advantages of coupling expression-level with functional analysis when it comes to fully understanding alternative splicing regulation[10,11]. To this end, we decided to complete our long read-defined transcript models with isoform-level functional information using IsoAnnotLite (https://isoannot.tappas.org/isoannot-lite/). IsoAnnotLite can incorporate functional information from sequence-based predictors and biological databases into any long read-defined transcriptome, as long as an already-available, compatible functional annotation -i.e. the ones available within the tappAS framework[10]- is provided. The set of compatible functional features (such as domains, motifs or binding regions for other molecular elements) is diverse, and encompasses both the transcript and protein isoform levels. All in all, we aimed to use the co-expression analysis pipeline in the *acorde* R package (https://github.com/ConesaLab/acorde) to predict the changes in inclusion of functional features encoded by isoform-level co-expression across cell types, and therefore the potential of co-selected alternative isoforms to exert different biological functions.

Briefly, IsoAnnotLite requires four inputs for functional annotation: SQANTI3 outputs **classification.txt*, **junctions.txt* and **corrected.gtf* files, each of them containing information regarding long read-defined transcripts, and a tappAS-compatible GFF3 file containing functional annotations. Functional features are then transferred from the GFF3 source annotation file to the GTF target annotation file. We run IsoAnnotLite v2.4 using SQANTI3 output files for our transcriptome, and a functional annotation GFF3 file generated after combining the mouse RefSeq functional annotation available in tappAS (RefSeq78), and a comprehensively-annotated mouse neural transcriptome belonging to neural progenitors[10]. We selected several functional categories from each of them, as detailed in Table 1. Once the input files are provided, IsoAnnotLite performs what we refer to as positional transfer of features. First, functional information that is positionally defined within isoforms in the GFF3 source file is translated to its corresponding genomic position. Then, functional feature information is mapped to the unannotated isoforms in the target GTF if the feature's genomic positions are included within any of the transcript's exons. The process is performed for both transcript and protein-level annotations, as well as for genomic functional information such as Gene Ontology (GO) terms, KEGG pathway information, if present. We run IsoAnnotLite (v2.6) using the following non-default parameters: *-gff3* (provides the combined GFF3 annotation file) and *-novel.*

As a result, 35028 isoforms in the GTF target annotation file (94.71% of 36986 isoforms in our transcriptome) were annotated, meaning that at least one functional feature could be successfully transferred from the source annotation file. The remaining 696 isoforms received no functional information due to the fact that the corresponding gene was not found in the GFF3 file, however, these represent but 1.88% of the long read-defined isoforms, meaning that feature transference was effective even though the reference GFF3 annotation files used did not exactly correspond to the same mouse samples, or even the same reference transcriptome version, namely RefSeq96 was used as the reference annotation for transcriptome construction and quality control using PacBio ENCODE data, while tappAS' mouse GFF3 functional annotation belonged to RefSeq version 78.

Regarding the comprehensiveness of the annotation process, IsoAnnotLite reported that ~47% of transcripts in the target GTF had the same associated reference transcript as one of the isoforms in the GFF3 reference annotation, which meant total agreement between their splice junctions and therefore ensured perfect possitional transference of features in the case of these isoforms. Even more reassuringly, feature transference in the 3’ and 5’ UTRs (regions that may differ even when the reference associated transcript is the same) is completed by IsoAnnotLite’s algorithm if the alternative UTR regions are included in other reference GFF3 transcripts, and contain functional features. For the remaining long read-defined transcripts, transference depended on the differences among alternative isoforms from the same genes present in the target and reference functional annotations, and was therefore expected to vary across genes. Consequently, the total number of functional features recovered was diverse depending on the functional category under evaluation (Table 1). It should be noted, however, that these numbers additionally rely on the extent to which these categories were originally present in the reference functional annotation files used, with less frequently annotated categories being often rare in their source GFF3 file (for instance, intramembrane protein regions). A complementary explanation has to do with the fact that the mouse NSC time series transcriptome was considerably more condition-specific than RefSeq, decreasing the possilibty to match the genes and isoforms included in the target GTF, and making transference challenging for categories extracted from it, such as RBPs and NLS.

| Source functional database | Feature category | No. of annotated functional features | Source GFF3 annotation file |
|---|---|---|---|
| *COILS* | COILED | 1184 | Mouse NSC time series |
| *CORUM* | Complex | 197 | Mouse NSC time series |
| *GeneOntology* | C | 32163 | Mouse RefSeq78 |
| *GeneOntology* | F | 30144 | Mouse RefSeq78 |
| *GeneOntology* | P | 30245 | Mouse RefSeq78 |
| *KEGG* | Pathway | 2652 | Mouse RefSeq78 |
| *MetaCyc* | Pathway | 1823 | Mouse RefSeq78 |
| *miRWalk* | miRNA binding | 186755 | Mouse RefSeq78 |
| *MOBIDB LITE* | DISORDER | 2732 | Mouse NSC time series |
| *NLS mapper* | MOTIF | 1020 | Mouse NSC time series |
| *NMD prediction* | NMD | 268 | Mouse RefSeq78 |
| *PAR-clip data* | RBP binding | 3000 | Mouse NSC time series |
| *PFAM* | CLAN | 23526 | Mouse RefSeq78 |
| *PFAM* | DOMAIN | 50736 | Mouse RefSeq78 |
| *Reactome* | Pathway | 11988 | Mouse RefSeq78 |
| *RepeatMasker* | Repeat | 35075 | Mouse RefSeq78 |
| *scanForMotifs* | 3UTRmotif | 4078 | Mouse NSC time series |
| *SIGNALP EUK* | SIGNAL PEPTIDE | 2641 | Mouse NSC time series |
| *TMHMM* | TRANSMEMBRANE | 1379 | Mouse NSC time series |
| *UniProtKB/Swiss-Prot* | ACTIVE SITE | 4790 | Mouse NSC time series |
| *UniProtKB/Swiss-Prot* | BINDING | 32126 | Mouse NSC time series |
| *UniProtKB/Swiss-Prot* | COILED | 3201 | Mouse NSC time series |
| *UniProtKB/Swiss-Prot* | COMPBIAS | 6392 | Mouse NSC time series |
| *UniProtKB/Swiss-Prot* | INTRAMEMBRANE | 385 | Mouse NSC time series |
| *UniProtKB/Swiss-Prot* | MOTIF | 15126 | Mouse NSC time series |
| *UniProtKB/Swiss-Prot* | PTM | 237128 | Mouse NSC time series |
| *UniProtKB/Swiss-Prot* | TRANSMEMBRANE | 19293 | Mouse NSC time series |
| *UTRsite* | 3UTRmotif | 44013 | Mouse RefSeq78 |
| *UTRsite* | 5UTRmotif | 623 | Mouse RefSeq78 |
| *UTRsite* | PAS | 8787 | Mouse RefSeq78 |
| *UTRsite* | uORF | 13370 | Mouse RefSeq78 |

**Table 1: Summary of IsoAnnotLite functional feature transference.** *Source functional database* refers to the original source of the functional information, which can be either a biological database or a sequence-based prediction tool. *Feature categories* are broad terms under which multiple entries of functional features encoding similar functions are classified. *Functional features* are the specific motifs, domains, regions, etc. annotated within the transcriptome's isoforms. *Source GFF3 annotation file* refers to the functionally annotated transcriptome from

which the feature information for each of the functional categories was obtained, and then passed on to IsoAnnotLite for feature transference. *NLS: nuclear localization signal; NMD: nonsense-mediated decay; NSC: Neural Stem Cells; PTM: post-translational modification; PAS: polyadenylation signal; RBP: RNA binding protein; uORF: upstream open reading frame.*
